# Alveolar Type II Cell-derived MMP1^high^ basal cells promote destructive microcysts in idiopathic pulmonary fibrosis

**DOI:** 10.64898/2025.12.08.693033

**Authors:** Kiana Garakani, Alexis N. Brumwell, Marylene Toigo, Anjali Jacob, Ying Wei, Tsung Che Ho, Paul J. Wolters, Michael A. Matthay, Regan Volk, Balyn Zaro, Stefano Iantorno, Asres Mitke, Dean Sheppard, Vincent Auyeung, Claude Jourdan Le Saux, Harold A. Chapman

## Abstract

Idiopathic Pulmonary Fibrosis (IPF) is a fatal lung disease characterized by progressive epithelial metaplasia and widespread fibrosis. Alveolar microcysts develop near terminal airways in IPF and are linked to poor outcome. Using HTII-280 as a short-term lineage marker of AT2-derived AT0 (SFTPC^+^/SCGB3A2^+^) and basaloid (KRT17⁺) cells, together with organoids and spatial transcriptomics (Xenium), we highlight epithelial similarities between respiratory bronchioles (RBs) and alveolar microcysts both having AT0, SCGB3A2+, and basaloid/basal cells (BCs), albeit with expanded BCs in IPF microcysts. The AT0- and AT2-derived BCs strongly express the collagenase, matrix metalloproteinase protein-1, MMP1 in organoids — mirroring *in situ* BCs lining IPF microcysts, but distinct from MMP1^low^ BCs in large airways or normal lungs. Expression of MMP1 correlates with basal cell hypoxia pathway activity.

MMP1^high^ AT2-derived BCs and IPF BCs promoted type 1 collagen degradation *ex vivo* and *in vivo* after xenotransplantation, forming microcystic structures that were abrogated by concurrent MMP inhibitor treatment. Notably, a Frizzled 5 WNT agonist antibody reversed the MMP1^high^ state of AT2-derived BCs, raising a possible therapeutic approach. These findings suggest AT2 transdifferentiation to basaloid/basal cells is uncommon in normal lungs but can expand as a potential source of alveolar destruction, likely contributing to the pernicious course of IPF.

## Introduction

Idiopathic Pulmonary Fibrosis (IPF) is a progressive disorder of aging characterized by widespread epithelial metaplasia with loss of normal AT2 and AT1 alveolar lining cells and progressive extracellular matrix (ECM) accumulation ultimately leading to respiratory failure (1–5). The clinical impact of limited treatments for fibrosis, especially for the ∼200,000 annual IPF cases, is significant. A key element in IPF pathology is the presence of subpleural foci of collagen-producing fibroblasts overlaid by dysfunctional epithelium and micro-cystic foci juxtaposed with near normal-appearing alveoli (usual interstitial pneumonia, UIP pattern) (6, 7). The extent of the UIP pattern correlates strongly with IPF outcome and poor treatment response (8, 9). Apparent coalescence of microcystic lesions over time, possibly prompted by subpleural mechanical strain, leads to the characteristic radiographic macro-honeycombing that defines the illness as UIP by CT imaging (10, 11). Moreover, UIP can develop in other chronic lung diseases such as hypersensitivity pneumonitis or autoimmune interstitial lung disease and again the appearance of UIP pattern portends a difficult to treat process (4, 7). How microcystic lesions are generated and mechanistically linked to underlying alveolar epithelial metaplasia and fibrosis is unclear.

The destructive nature of IPF pathobiology suggests an important role for ECM-degrading proteases. Proteases, and in particular metalloproteases (MMPs), have been implicated in the pathobiology of pulmonary fibrosis for decades (12). Several MMPs have been linked to ECM degradation and pro-fibrotic signaling. Yet, because most studies are correlative or in mice—and selective inhibitors are unavailable — causal roles for individual MMPs in humans are unresolved. Nonetheless, the MMP most consistently associated with pulmonary fibrosis and progression of IPF in particular is elevated blood levels of MMP7(13). In addition, MMP1 is most elevated in IPF and found together with MMP7 to discriminate IPF from other ILDs (14).

MMP7 mainly cleave basement membrane collagen IV, along with other non-collagen matrix proteins such as fibronectin. MMP1 is the major adventitial collagenase of type 1 and 3 fibrillar collagens accumulating in IPF, though its role in lung injury or fibrosis is not well defined(15, 16). We previously reported that MMP1 and MMP7 mRNA are in the top 25 of genes upregulated in both IPF and AT2-derived basal cells (17). Analysis of IPF by spatial transcriptomics revealed that a subset of IPF alveolar basal cells account almost entirely for the increased expression of MMP1 in IPF. We therefore undertook a more detailed study of the location and function of MMP1^high^ alveolar basal cells.

## Results

### HTII-280 is a short-term lineage trace for KRT17+ cells and bronchiolized IPF microcyst epithelium

The recognition of HTII-280 as an empiric antigen epitope for human AT2 cells has been known for decades and used to isolate this cell population in many prior studies by flow cytometry (5, 18, 19). We inspected the distribution of HTII-280 immunostaining in both normal and IPF lung sections. Consistent with prior reports indicating the normal respiratory bronchioles (RBs) are comprised in part by SFTPC^+^/SCGB3A2^+^ cells (termed AT0), we found HTII-280^+^ AT0 cells in virtually all RBs in five normal lung sections examined (4-5 RBs per section) (Figure 1A) (20). RBs also contained numerous HTII-280^neg^ SCGB3A2^+^ cells as expected, and rare basal cells (KRT5^+^). No normal appearing RBs were found in IPF lung sections. Surprisingly, microcystic structures in IPF were comprised of widespread HTII-280 stained epithelium, with foci of AT0 and SFTPC^neg^/SCGB3A2^+^ cells as well as larger areas of basal cells and other airway-like cells that were HTII-280+ (Figure 1A). The larger areas of the open cysts lined by airway-like cells are consistent with the term bronchiolization applied to this cytopathology for decades, interestingly the frequent co-staining with HTII-280 raised the question of their origin. To further investigate this point, we capitalized on recent findings identifying the source of HTII-280 antibody binding. A HTII-280 immunoprecipitate of whole human lung lysate was examined by LC/MS and the top “hit” for membrane-linked protein in this analysis was MUC1 (Supplemental Table 1). HTII-280 immunoprecipitates were recognized by MUC1 antibody and vice versa (Figure 1B). The protein smear in the immunoprecipitate suggested extensive glycosylation. Indeed, all HTII-280 staining of lung lysates was abolished by 1-hour deglycosylation consistent with the presence of glycosyl epitopes in antibody binding uniquely identifying AT2 cells (Figure 1C). Likewise, binding of HTII-280 to HTII-280-flow sorted normal AT2 cells was transiently abolished by short term exposure to neuraminidase but reappears in ∼12 hours of culture consistent with the high expression level of MUC1 and the HTII-280 epitope (Supplemental Figure 1D).

**Figure 1.**
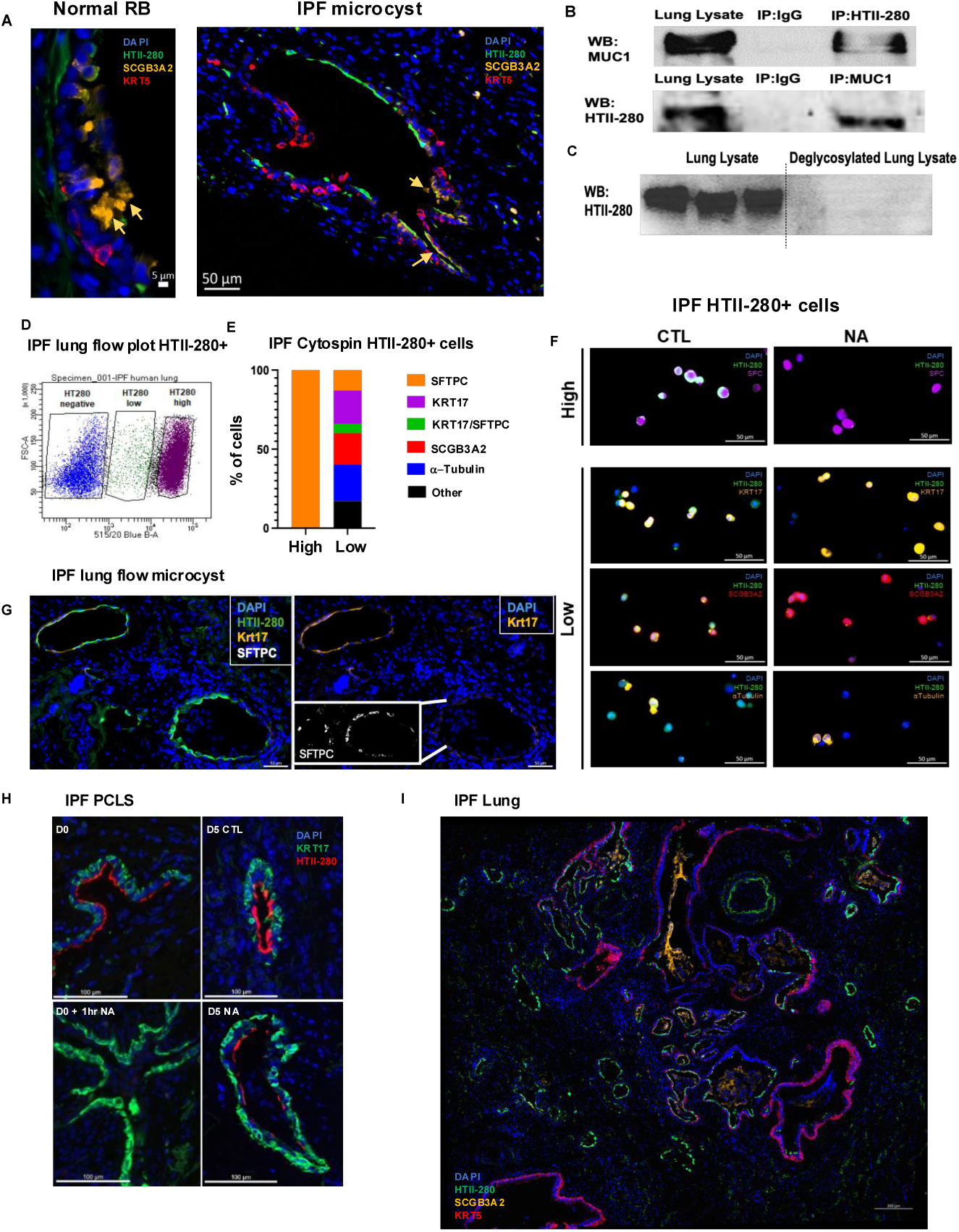
HTII-280 is a lineage marker for AT2 differentiation and microcystic structures in IPF are derived from AT2. (**A**) Representative image of normal respiratory bronchiole from healthy donor (left) and microcystic structure from IPF lungs (right) stained by immunofluorescence (IF) for HTII-280 (green), SCGB3A2 (yellow), and KRT5 (red). Yellow arrows indicate SCGB3A2^+^ cells. (**B**) Western blot for MUC1 and HTII-280 on lung lysate and immunoprecipitates. (**C**) Western blot for MUC1 from lung lysate with or without prior deglycolysation with neuraminidase (NA) and PNGaseF and O-deglycosidases. (**D**) Representative flow blot of sorted IPF HTII-280+ cells indicative of two distinctive populations, HTII-280^low^ and HTII-280^hi^ cells from an IPF lung above the MFI of the isotype control (Supplemental Figure 1). Similar flow plots were found in >10 biological replicates. (**E**) Cell-type distribution (percent) from cytospins of freshly sorted IPF HTII-280^hi^ and HTII-280^low^ stained by IF for KRT17, SCGB3A2, α-TUBULIN, and SFTPC; n = 100–150 cells per condition. (**F**) Representative images of HTII-280^hi^ and HTII-280^low^ cytospins ± NA, stained by IF for HTII-280 (green), SFTPC (purple), KRT17 (yellow), SCGB3A2 (red), or α-TUBULIN (yellow). (**G**) Representative image of microcystic structure from IPF lung stained by IF for HTII-280 (green), KRT17 (yellow) and SFTPC (white). (**H**) Representative image of PCLS from IPF lungs containing microcystic structure treated for one hour with NA, collected immediately and 5 days later and stained for KRT17 (green) and HTII-280 (red). Similar findings were observed in 3 biological replicates. (**I**) Distribution of HTII-280 immunostaining in IPF lung section stained for HTII-280 (green), SCGB3A2 (yellow), and KRT5 (red). Similar findings were observed in four separate IPF lung explants.

As previously reported, HTII-280 flow plots of epithelial cells recovered from normal lungs are comprised of a large HTII-280 high (HTII-280^hi^) cell cluster and a smear of lower intensity staining we term HTII-280^low^, but above the mean fluorescent intensity (MFI) of the isotype control (Figure 1D, Supplemental Figure 1A, B). In a normal lung, both HTII-280^+^ high and low cells expressed SFTPC (Supplement Figure 1C). However, immunostaining of HTII-280-sorted cells from IPF lungs revealed that the HTII-280^hi^ population was virtually entirely SFTPC+ whereas the HTII-280^low^ population consisted of a heterogenous cell population some of which were SFTPC^+^, SCGB3A2^+^, α-TUBULIN^+^, KRT17^+^, or KRT17^+^/SFTPC^+^ (Figure 1E, F). There was also a HTII-280^neg^ population not further characterized of similar MFI as the isotype control. Treatment of HTII-280^low^ cytospins with neuraminidase prior to fixation abolished HTII-280 staining in all of the airway lineages (Figure 1F), consistent with their common derivation from HTII-280^+^ AT2 cells by transdifferentiation as previously reported in organoid co-cultures (17, 21). Furthermore, HTII-280 was found in cells expressing KRT17 in several microcystic structures in IPF, consistent with their common derivation from AT2 cells by transdifferentiation, as previously reported in organoid co-cultures and PCLS (Figure 1G) (21, 22). To further test this point, human precision-cut lung slices (PCLS) freshly prepared from IPF lungs were treated with neuraminidase and followed over time in culture up to 5 days (Figure 1H). Neuraminidase abolished the HTII-280 signal in KRT17^+^ in IPF PCLS after one hour. New signal was first appreciated after three days and by five days was comparable to sections never exposed to neuraminidase. It was notable that not all KRT17^+^ cells stained with HTII-280 in fresh IPF lung sections and this was variable among sections. An example of KRT17^+^/SFTPC^neg^/HTII-280^+^ cells contrasted with KRT17^+^/SFTPC^+^/HTII-280^+^ cells is illustrated in Figure 1G. Two IPF microcysts are present in the IPF section, one which is clearly SFTPC^+^ (lower) and one which is SFTPC^neg^ (Figure G, upper). Both microcysts are stained by HTII-280 though the intensity of staining is less in the Krt17^+^ cells lining the upper cyst, consistent with HTII-280-sorted cells from IPF lungs. To extend this line of experiments more directly to IPF cytopathology, frozen and fixed IPF alveolar region sections were immunostained for HTII-280. Pseudo-stratified large airways were uniformly HTII-280 negative whereas microcystic structures near the airway termini were largely HTII-280 positive, including some basal cells (Figure 1I). These findings suggest many if not most IPF microcysts, like RBs, are lined by AT2-derived airway lineages.

### Comparison of RB and microcyst epithelial populations: identification of MMP1^hi^ basal cells

To further explore the epithelial composition of RBs and IPF microcysts, we turned to spatial transcriptomics (Xenium) for higher resolution. The patient characteristics are described in Supplemental Table 2. Respiratory bronchioles in normal lungs were identified by linear epithelial arrays near the end of bronchioles and surrounded by AT2 alveoli and annotated by canonical markers for RB epithelium: AT2 (ABCA3^+^), AT0 (ABCA3^+^/SCGB3A2^+^), and ABCA3^neg^/SCGB3A2^+^ cells (Figure 2A). Multiple sections were analyzed and the epithelial composition quantified (Figure 2A right). As expected, the main components of the RBs were ABCA3^neg^/SCGB3A2^+^ (60.6 +/-18.1%) and ABCA3^+^/SCGB3A2^+^(24.4 +/- 11.8%). Very few cells expressed KRT17 (6.1 +/- 4.4%) or KRT5 (1.6 +/- 2.1%). In contrast, the cell distribution in IPF microcysts was very distinct (Figure 2B). IPF microcysts were identified as open structures < 2mm in longest dimension, with an epithelial lining layer, and unattached to an obvious small airway. Cell identities quantified in Figure 2B were based on canonical markers for cell types identified in Figure 2A. ABCA3 was used as a surrogate AT2 marker for SFTPC in this analysis because of better performance in the spatial transcriptomic Xenium arrays. The epithelial cell types were similar between RBs and IPF microcysts except for marked expansion of KRT17^+^/KRT5^neg^ (ABI2s a.k.a. aberrant basaloid cells) accounting for 27.3 +/- 10.3 % of the cell population and KRT5^+^ basal like cells (46.9 +/- 16.8 %) as indicated in Figure 2B. Inspection of IPF microcysts consistently identified MMP1 in microcyst basal cells (Figure 2C, MMP1 transcripts in white). Other common cells in these cysts such as ABI2s, ABCA3± SCGB3A2 were MMP1 negative. A UMAP representation derived from Xenium analysis of all cell types in IPF and normal donor lungs revealed MMP1 expression exclusively in IPF epithelial KRT5^+^ basal cells (Figure 2D and Supplemental Figure 2A). Spatial transcriptomic imaging indicated that virtually all microcyst basal cells express MMP1. However, low levels of MMP1 transcript were detectable sporadically in basal cells throughout conducting airways. Feature plots of MMP1 expression in basal cell subsets of single-cell RNA sequencing (scRNA seq) files confirmed little or no MMP1 in normal lung basal cells but clear upregulation in IPF and organoid-derived basal cells (Figure 2E). Differential expressed genes in IPF basal cells compared to normal lung basal cells confirmed the upregulation of MMP1 expression in IPF basal cells (Supplemental Table 3). MMP1 is reportedly induced by hypoxia (23). Indeed, subsetting MMP1^hi^ and MMP1^negative^ basal cell scRNA seq indicated higher hypoxia pathway activity in the MMP1^hi^ population (Supplemental Figure 2B). Overall, these findings indicate basal cell MMP1 levels are high in IPF alveolar-derived basal cells, largely in microcysts, and raise the possibility that fibrillar collagen destruction, prominent in IPF, is mediated in part by basal cell release of interstitial collagenase MMP1. It should be noted that not all microcysts are lined by basaloid/basal cells. Consistent with a previous Xenium analysis of fibrotic lungs (24), some cyst-like structures were lined by AT2 cells, some of which also contained SCGB3A2+ cells. Their origin is unclear and not explored in this manuscript.

**Figure 2.**
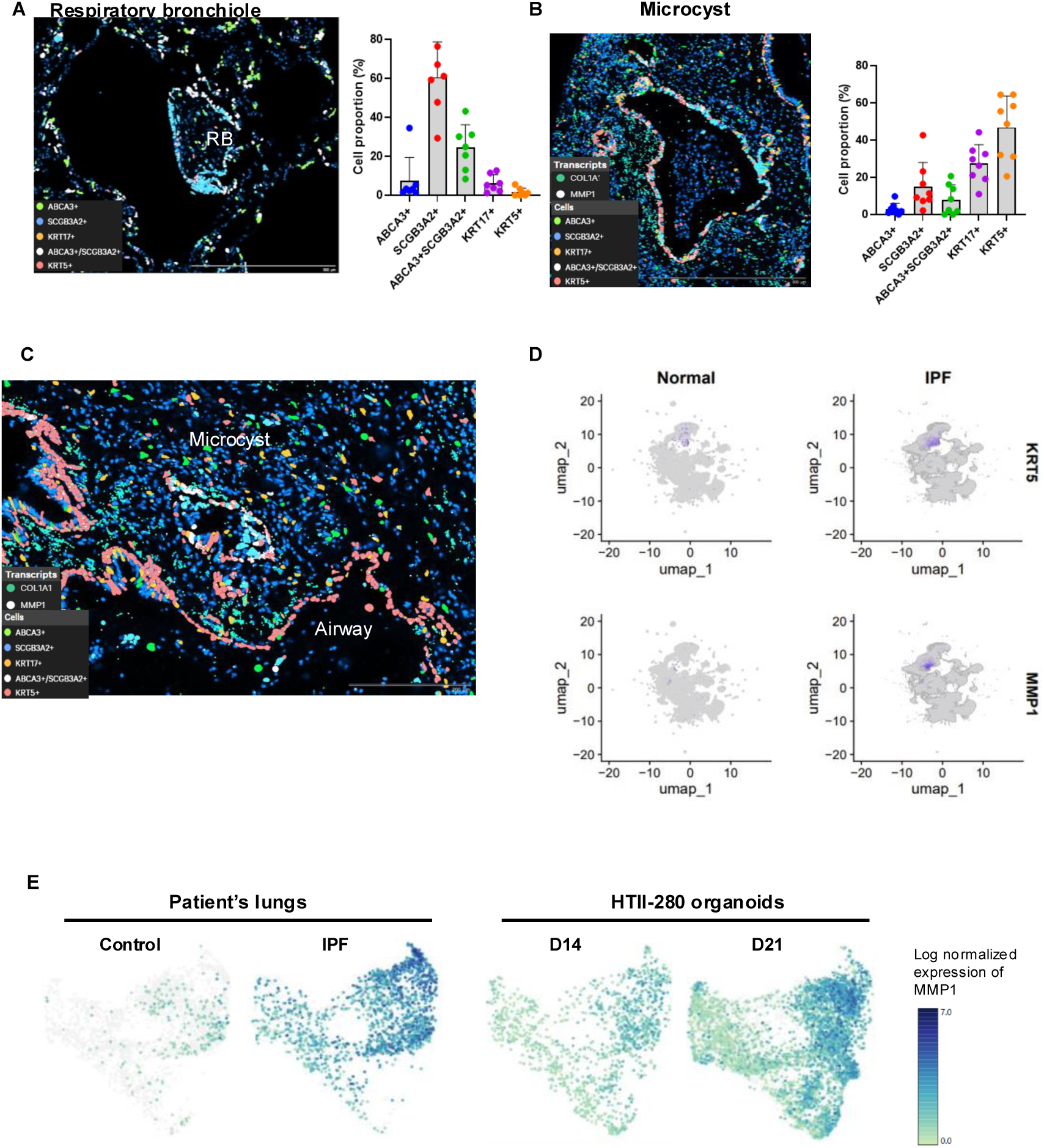
Microcysts in IPF lungs contain mostly basal cells, expressing a high level of MMP1. (**A**) Respiratory bronchiole (RB) identified in normal lung sections by a short linear sequence of airway epithelium surrounded by alveoli. Slides hybridized with Xenium probe set and annotated for cell types using markers shown on the image. Average composition of cell types quantified with each dot representing a distinct RB. (**B**) Representative microcyst identified as <1mm diameter open epithelial-lined cyst and annotated for cell type by canonical markers indicated on the image. Cell composition quantified with Xenium explorer after cell type annotation on R. MMP1 transcript (white) localizes to basal cells (red) and preferentially to basal cells overlying COL1A1 (green) fibroblasts. (**C**) MMP1^hi^ microcystic structure and MMP1^low^ airway from IPF lungs. (**D**) KRT5 and MMP1 expression in UMAP display of single-cell RNA sequencing control and IPF human lung from spatial transcriptomic probe set (n = 7 IPF and 3 control biological replicates). (**E**) MMP1 expression in single-cell RNA sequencing of control, IPF lung, and AT2-derived basal cells from organoids after 14 and 21 days of culture.

We previously reported that CEACAM6^+^/SCGB3A2^+^ cells, isolated by flow cytometry, differentiated to AT2 cells when placed in an AT2-promoting culture medium (25). However, when we repeated the isolation of CEACAM6^+^/HTII-280^neg^/NGFR^neg^ RB cells isolated by flow cytometry from normal lungs and co-cultured them with adult human lung fibroblasts in organoids for 9 days, virtually all the colonies were strongly KRT17^+^/KRT5^+^ basal cells (Figure 3A-E; (17)). Organoids of NGFR^+^ basal cells from the same sort are shown for comparison (Figure 3C). In addition, we confirmed by gene expression, that the EPCAM^+^ cells isolated from the 9-day CEACAM6 and basal cells derived organoids expressed high levels of *KRT5, KRT17*, and *MMP1* mRNA and decreased levels of *SCG3A2* mRNA (Figure 3E). This result does not mean there are necessarily multipotent progenitors within normal RBs, but the population of cells at least contains progenitors of both AT2 or basal-like cells when placed in differentiating conditions.

**Figure 3.**
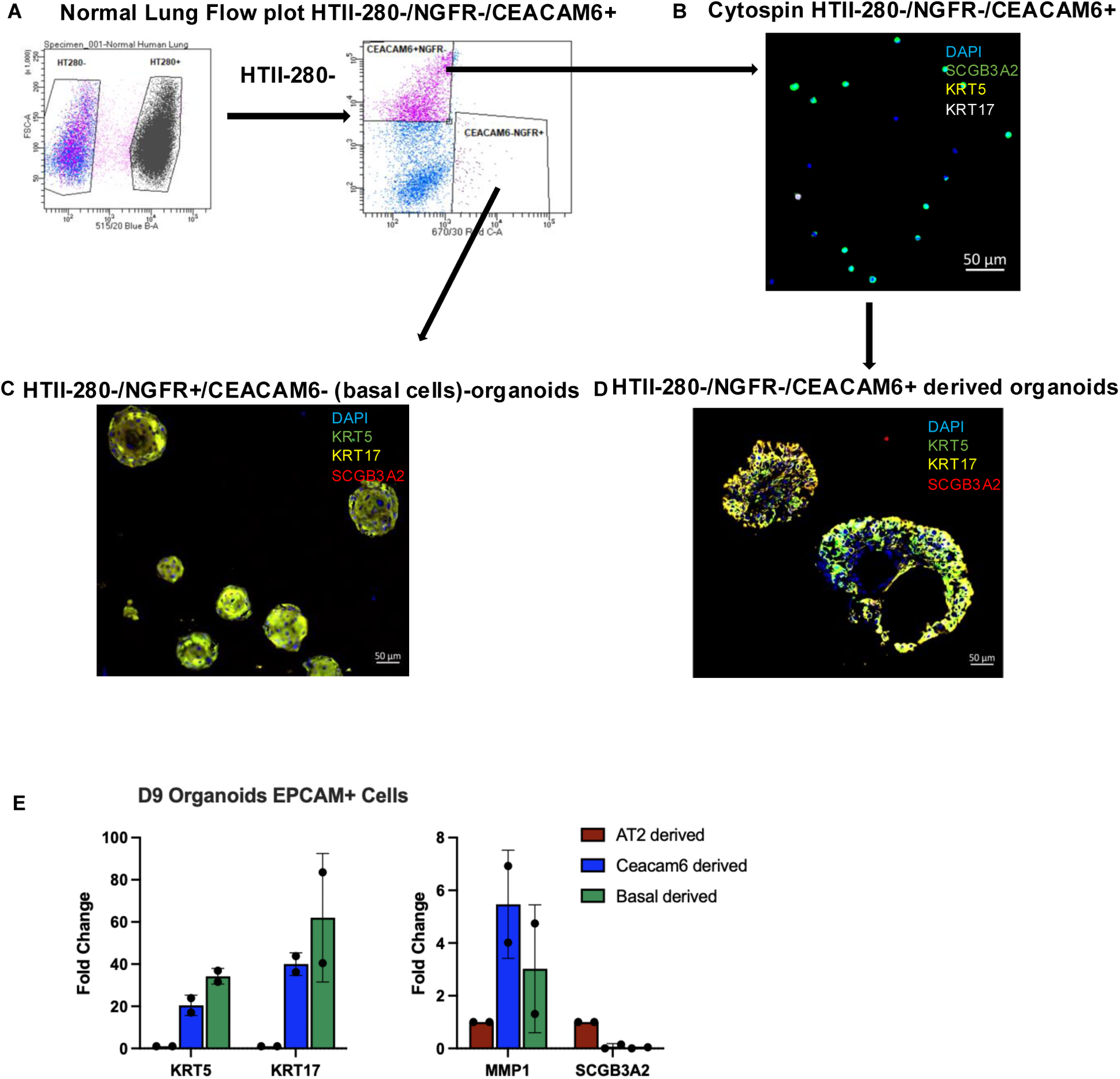
AT2 and RB cells transdifferentiation to MMP1hi basal cells. (**A**) Representative flow plot of sorted HTII-280^-^ cells (left), and CEACAM6^+^NGFR^-^, and CEACAM6^neg^NGFR^+^ cells from normal human lungs by FACS (right). Immunostaining confirmed CEACAM^+^NGFR^-^cells cells were <1% basal. (**B**) Representative image of HTII-280^neg^CEACAM^+^NGFR^neg^ cytospin, stained by IF for SCGB3A2 (green), KRT5 (yellow), or KRT17 (white). (**C, D**) IF of basal cell-derived (**C**) and Ceacam cell-derived organoids with primary fibroblasts organoids after 9 days of culture staining for KRT5 (green), KRT17 (yellow), SCGB3A2 (red) (n = 2). (**E**) *KRT5, MMP1* and *SCGB3A2* mRNA relative expression of epithelial cells (EPCAM+) sorted from basal or Ceacam-derived organoids after 14 days of culture. Relative change in mRNA expression is presented as fold change normalized to the EPCAM^+^ fraction of 9-day AT2-derived organoids (n = 2).

### Basal cell MMP1 mediates Type 1 collagen degradation both in vitro and in vivo

To test whether AT2 derived basal cells not only express *MMP1* mRNA (Figure 2E) but also degrade collagen, we cultured 21-day AT2-derived basal cells on immobilized type 1 collagen for 8 days with or without MMP inhibitor treatment (GM6001). Conditioned media were collected and incubated with DQ-collagen I for an hour. Soluble fluorescent collagen fragments were measurably released but became undetectable when cells were treated with GM6001 (Figure 4A). To test the impact of basal cell MMP1 *in vivo*, we next turned to xenotransplantation.

**Figure 4.**
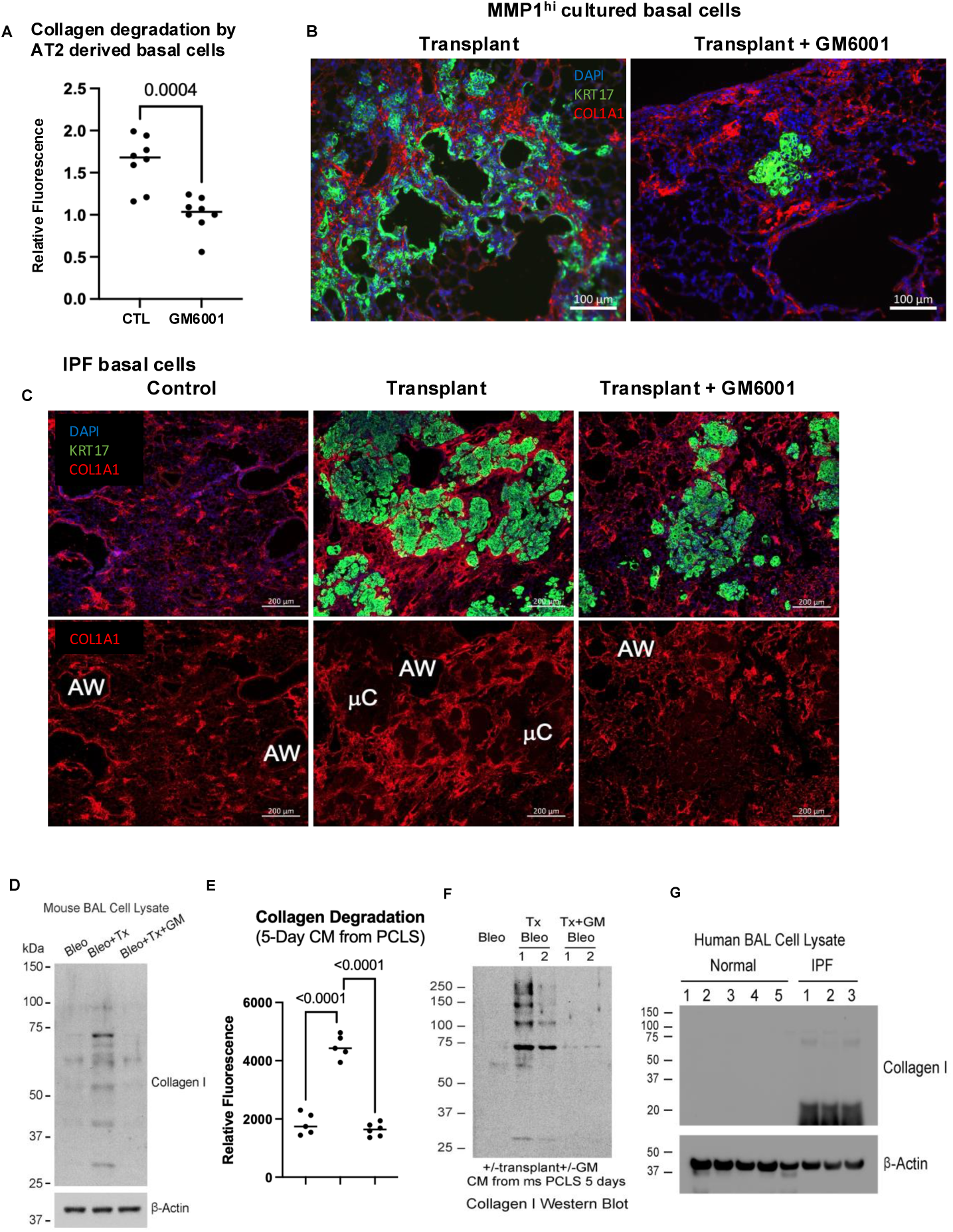
Human MMP1^hi^ basal cells degrade type I collagen and form cyst in vivo, both suppressed by GM6001. (**A**) Fluorescent collagen degradation assay on conditioned media from 21-day organoid AT2-derived basal cell after 8 days of culture treated with or without GM6001 (n =3). (**B**) Representative image of human MMP1^hi^ basal cells transplanted into bleomycin injured NSG mice treated with or without GM6001 stained for human-specific KRT17 (green) and COL1A1 (red). (**C**) Representative lung section images of bleomycin NSG mice transplanted with IPF BCs, treated with or without GM6001 stained for human-specific KRT17 (green) and COL1A1 (red) (AW = Airway, μC = Microcyst). (**D**) Western blot of degraded type I collagen in BAL cell lysates from bleomycin NSG mice transplanted (Tx) with IPF BCs, treated with or without GM6001. (**E**) Fluorescent collagen degradation assay on 5-day conditioned media of PCLS from lungs of bleomycin NSG mice transplanted with IPF BCs, treated with or without GM6001 (n = 4 – 6, 2 independent cohorts). (**F**) Western blot of degraded type I collagen in concentrated conditioned media (CM) of PCLS from lungs of bleomycin NSG mice transplanted (Tx) with IPF BCs, treated with or without GM6001. (**G**) Western blot of degraded type I collagen from control and IPF BAL cell lysates (n = 5 normal donor and 3 IPF). Magnification of an airway and microcyst characterized by the non-presence of cells. Statistical significance was determined by unpaired t-test (**A**) and Dunnett’s multiple comparisons test (**E**).

Ten days after transplant, MMP1^hi^ basal cells, identified by human-specific KRT17 antibody, engrafted in the bleomycin injured regions of NOD.Cg-*Prkdc^scid^ Il2rg^tm1Wjl^*/SzJ (NSG) mouse lungs and formed microcystic-like structures lined by KRT17+ cells (Figure 4B); these structures did not form in mice receiving GM6001. We next examined freshly sorted IPF basal cells in xenotransplants. Isolated cells were transduced by lentivirus with a mCherry reporter to facilitate localization of transplants. After 10 days post-transplant, collagen degradation beneath KRT17^+^ IPF basal cells coincided with formation of collagen1-depleted zones resembling microcysts (Figure 4C). Additional sectioning of these transplants and higher magnification revealed cyst-like areas found in all IPF basal transplants. Treatment with GM6001 abrogated both collagen depletion and cystic zones (Supplemental Figures 3E, F). To verify collagen degradation *in vivo* after xenotransplantation, we capitalized on the capacity of resident alveolar macrophages to internalize and degrade partially degraded collagens (26, 27). Twenty days after bleomycin injury corresponding to 10 days after transplant, collagen fragments were barely detected in the broncho alveolar lavage (BAL) from bleomycin injured mice but clearly increased in mice that were transplanted with IPF basal cell (Figure 4D, Supplementary Figure 4A, B).

Treatment with GM6001 virtually abrogated the increased collagen fragments in IPF basal cell transplants (Figure 4D, Supplemental Figure 4A, B). To further link IPF basal-cell transplantation to collagen degradation and microcyst formation *in vivo*, we prepared PCLS containing engrafted cell areas identified as mCherry^+^ from bleomycin injured, engrafted mice ± GM6001. The conditioned media concentrated from these PCLS showed increased collagen-degradation activity in transplanted samples (fluorescent collagen degradation assay, Figure 4E) and increased solubilized total collagen fragments by Western blot (Figure 4F, Supplementary Figure 4C, D) compared to bleomycin controls and PCLS of transplanted and GM6001 treated mice. In all assays, collagen degradation was abolished when the transplanted mice were also treated with GM6001.

Together, these results show that transplanted IPF basal cells drive collagen degradation and microcyst-like structure formation, and that inhibiting their MMP activity with GM6001 prevents both. Finally, we asked whether resident alveolar macrophages in human IPF lungs contained type 1 collagen fragments. Whereas BAL cell pellets of normal control samples had no detectable intracellular collagen fragments, all of the IPF samples were strongly positive for partially degraded type 1 collagen (Figure 4G).

### Simultaneous canonical WNT activation and suppression of non-canonical WNT blocks and can reverse AT2 to basal cells transdifferentiation and MMP1 expression

We previously reported that AT2 to basal cells transdifferentiation is driven by non-canonical WNT signaling through the interaction of sFRP2 with Frizzled-5 (Fzd5) (21). A frizzled-5 agonist (Fzd5Ag) has been shown to stimulate alveolar epithelial stem cell activity by promoting the activation of canonical WNT pathway (28). Consistent with this, in our organoid system, Fzd5Ag increased *AXIN* 2 mRNA expression in the EPCAM+ cells (Figure 5A). Fzd5Ag inhibited sFRP2-induced nuclear accumulation of NFATc3 in HEK293 cells overexpressing FZD5 (Figure 5B).

**Figure 5.**
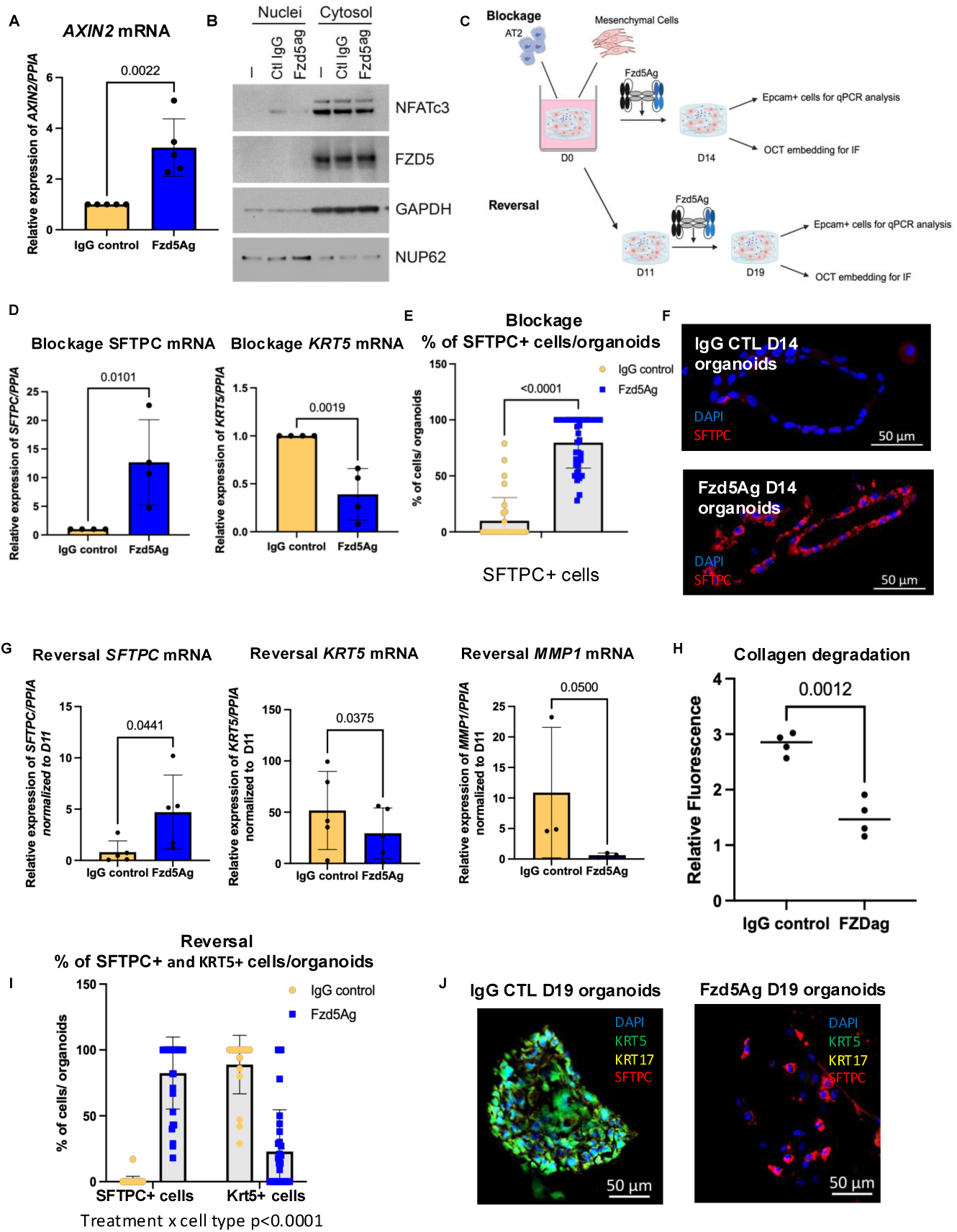
MMP1hi basal cell state is prevented and can be reversed by Frizzled 5 agonist. (**A**) *AXIN2* mRNA relative expression of epithelial cells (EPCAM+) sorted from organoids treated with IgG control or Frizzled-5 agonist (Fzd5Ag) from day 1 to day 14 (n = 5). (**B**) Western blot for NFATc3, FZD5, GAPDH, and NUP62 on nuclei and cytosol of epithelial cells treated with sFRP2 as a Fzd5 ligand promoting nuclear NFATc3 and with IgG control or Fzd5 agonist (Fzd5Ag). (**C**) Experimental design of blockage and reversal assays with Fzd5Ag on organoids. (**D**) *SFTPC and KRT5* mRNA relative expression of epithelial cells (EPCAM+) sorted from organoids treated with IgG control or Fzd5Ag from day 1 to day 14 of culture (n = 3). (**E**) Percentage of SFTPC+ cells per organoids treated with IgG control or Fzd5Ag from day 1 to day 14 (n = 2 biological replicates). Each data point represents an organoid. (**F**) Representative image of IgG control or Fzd5Ag treated organoids stained by IF for SFTPC (red). (**G**) *SFTPC*, *KRT5*, *MMP1* mRNA relative expression of epithelial cells (EPCAM+) sorted from organoids treated with IgG control or Fzd5Ag from day 11 to day 19 (n = 3 – 4). (**H**) Type I collagen degradation assay of AT2-derived BCs treated with IgG control or Fzd5Ag from day 11 to day 19 of culture (n = 2 biological replicates and n = 2 technical replicates). (**I**) Percentage of SFTPC+ or KRT5+ cells per organoids treated with IgG control or Fzd5Ag from day 11 to day 19 of culture (n = 3 biological replicates). Each data point represents an organoid. (**J**) Representative image of organoids treated with IgG control of Fzd5Ag from day 11 to day 19 of culture stained by IF KRT5 (green), KRT17 (yellow), and SFTPC (red). Statistical significance was determined by unpaired t-test (**A, D**, and **H**), paired t-test (**G**), and mixed-effects analysis (**E** and **I**).

Hence, Fzd5Ag promotes canonical WNT signaling while suppressing noncanonical WNT signaling. To determine whether simultaneously promoting canonical WNT signaling and suppressing non-canonical WNT signaling will block the AT2 to BC transdifferentiation and/or promote reversal of AT2-derived BC to AT2, we treated AT2/mesenchymal cell organoids for 14 days starting on day 1 (blockage) or for 7 days starting on day 11 (reversal) (Figure 5C).

In 14-day human AT2 cells and human mesenchymal cells co-culture organoids treated with Fzd5Ag throughout the 14 days, *SFTPC* and *ABCA3* mRNA levels increased, accompanied by reduced *KRT5* mRNA expression, consistent with maintenance of AT2 identity rather than acquisition of basal cell fate transdifferentiation (Figure 5D and Supplemental Figure 5A). Cell-type composition analysis corroborated these findings: in IgG-treated controls, all organoids lacked SFTPC⁺ cells; in Fzd5Ag-treated cultures, all organoids contained SFTPC⁺ cells (Figure 5E, F).

We next asked whether Fzd5Ag treatment could reverse established transdifferentiation. Eleven-day AT2/Mesenchymal cell organoids were treated with Fzd5Ag for 7 days. Similar to the blockage experiment, Fzd5Ag increased expression of AT2 genes (*SFTPC*, *ABCA3*, *ETV5*) and decreased expression of basal cell genes (*KRT5*, *KRT17*, *TP63*, *MMP1*), indicating that activation of canonical WNT coupled with inhibition of noncanonical WNT enables AT2-derived basal cells to reacquire AT2 identity (Figure 5G and Supplemental Figure 5B). The decreased expression of *MMP1* mRNA induced by Fzd5Ag treatment was also associated with a decrease in collagen degradation capability of the treated cells (Figure 5H). Consistently, IgG-treated organoids were composed exclusively of KRT5⁺ cells, whereas Fzd5Ag-treated organoids were predominantly SFTPC⁺ with a small KRT5⁺ fraction (∼23%) (Figure 5I, J).

Collectively, these data indicate that Fzd5Ag both prevents and partially reverses AT2 to basal transdifferentiation while abrogating MMP1 expression, by shifting signaling toward canonical WNT and away from non-canonical WNT.

## Discussion

Accumulating evidence supports the paradigm that human AT2 cells are quite plastic and engaged not only in self-renewal but transdifferentiation to at least 3 distinct lineages: AT0 cells populating RB epithelium (20), AT1 cells spreading to create an alveolar epithelial barrier and support a vascular niche (29), and basaloid/basal cells in response to more severe alveolar injury (17). Studies reported here provide new details on AT2 plasticity as it relates to the origin of alveolar basaloid/basal cells and their distinct features in IPF cytopathology. IPF alveolar basal cells could arise from alveolar AT2 cells, RB epithelial progenitors as reported here (Figure 3), or by migration from terminal airways (or conceivably even larger airways) as is prominent in murine lungs responding to alveolar injury (30, 31). Our data favor the view that some and possibly all of the basal cells (and basaloid cells) arise from AT2 cells as judged by the widespread expression of low levels of HTII-280 epitope previously thought to be specific for AT2 cells on several airway lineages including ciliated cells, all known to be progeny of AT2-derived basal cells in organoids (17). That the sorting by flow cytometry recognizes progeny of AT2 cells is verified by the removal of the HTII-280 signal with neuraminidase. Although the HTII-280 protein antigen defined here as MUC1 is expressed widely in epithelial cells, the complex carbohydrate structure of AT2 cell MUC1 accounting for HTII-280 antibody binding appears to be unique. This may reflect the extensive glycosylation of AT2 cells as part of the maturation of surfactant protein C, highly expressed and unique to AT2 cells. Our finding that HTII-280 binding slowly decreases with time after transdifferentiation as judged by flow cytometry as well as epitope re-expression on KRT17^+^ basaloid cells after neuraminidase exposure supports an AT2 origin. Nonetheless, the conclusion that nearly all “bronchiolization” in fibrosis ultimately arises from AT2 transdifferentiation would be premature, requiring further exploration. Some IPF alveolar airway-like lineages are HTII-280^neg^, raising the possibility of multiple origins or reflecting their greater resident time in tissues after AT2 transdifferentiation.

We also could not exclude the possibility that some airway lineages that are commonly MUC1^+^ develop the AT2-glycosylyl program to generate the HTII-280 epitope in the context of IPF.

In contrast to most autoimmune interstitial lung diseases, the cytopathology of IPF leads to progressive tissue distortion and alveolar collapse/destruction from the earliest stages of diagnosis and this correlates with the expansion of airway-like epithelial lineages in the alveolar compartment over time, in part by replacing loss of AT2 cells from senescence and apoptosis but also loss of AT2 cells by their transdifferentiation to airway-like cells. It is also well known that destruction and distortion are caused by the fibrotic process itself, continued expansion of the ECM with progressive stiffness and vascular obliteration. Nonetheless, a strong component of this destructive process is the accumulation of microcystic areas that likely expand by collagen-rich ECM degradation as demonstrated here and coalesce over time into macro-cystic or honeycomb cysts discernible by chest CT. The extent of accumulation correlates with disease outcome (9). A surprising finding in our studies was the specific upregulation of the collagenase MMP1 in IPF basal cells lining most of the microcysts. Basal cell MMP1 upregulation in IPF is evident in scRNA seq files but its striking specificity is only revealed by spatial transcriptomic analysis (Xenium). This distribution provides a mechanistic explanation of why and how small basal cell clusters could use interstitial collagenase to evolve into larger, more destructive cysts, as we demonstrate directly in transplants of MMP1^hi^ basal cells into injured mouse lungs. Recent anatomical studies of IPF cytopathology reported loss of recognizable terminal bronchioles and the emergence of honeycomb cysts in the same or nearby space, consistent with a contribution from dysplastic respiratory bronchioles as shown here by the emergence of MMP1^hi^ basal cells from normal RB progenitors in organoids (32).

The position of RBs at the junction between terminal conducting bronchioles that contain basal cells and normal alveoli that do not, raises the likelihood that RBs are a human equivalent to the BASCs and other undifferentiated distal airway cells found in mice (30, 33). In that context, the basal cells in normal human terminal airways could arise from RBs, a possibility that bears exploration in future studies. Basaloid and basal cells also appear to have distinct functions.

Basaloid (KRT17^+^/KRT5^-^) cells emerge robustly in response to TGF-β1 signaling, exhibit several Epithelial-Mesenchymal Transition features, and engage in a set of fibroblast interactions after injury that could be reparative, along with AT1 differentiation, or pro-fibrotic if left unchecked. In contrast basal cells in the alveolar compartment outside of the airway tree are intrinsically pathological. Not only are the cells non-functional in gas exchange, RBs contain stem cell progenitors of numerous airway cell types that accumulate, and our data suggest that they promote, the disordered IPF cytopathology. This conclusion is fueled by the discovery reported here that alveolar basal cells accumulating in IPF, or other disorders with UIP pathology, express the fibrillar collagenase MMP1 and localize near collagen producing fibroblasts ((17), Figure 2).

Pharmacologic activation of canonical WNT signaling using Fzd5Ag also coincide with suppression of non-canonical WNT activity (Figure 5). This is mechanistically relevant given our prior work establishing that non-canonical WNT signaling is a key driver of AT2-to-basal transdifferentiation (21). Consistent with this model, the AT2-derived basal cells that emerge under disease-like conditions exhibit high MMP1 expression, a phenotype we associate with matrix degradation and the destructive microcystic pathology characteristic of IPF. Importantly, we show that this transdifferentiation program is not only preventable but reversible: enhancing canonical WNT while inhibiting non-canonical signaling halts the acquisition of a basal-like state including MMP1 expression. The relative contribution of canonical pathway promotion versus non-canonical pathway inhibition to this blockage and reversal remains to be dissected. Nevertheless, our data identify a tractable axis with immediate translational implications: Fzd5Ag or functionally equivalent strategies could preserve the AT2 pool in IPF and/or attenuate MMP1-high basal cells within microcystic structures, thereby limiting tissue destruction. Together, these results open a therapeutic avenue aimed at preventing further AT2 loss and mitigating protease-driven damage in IPF.

Our data also open another potential therapeutic avenue focusing on MMP1. Already MMP1 has been implicated in COPD and particularly emphysema development and progression, consistent with the destructive nature of MMP1^hi^ basal cells demonstrated here and with earlier studies by the D’amiento group (16, 34). However, a significant hurdle for development of drugs targeting MMP1 however is that no specific inhibitor of MMP1 activity (or that of most other MMPs) has been developed. Our finding that activation of/inhibition of non-canonical WNT signaling in AT2-derived basal cells reverses the MMP1^hi^ state invites additional studies focused on attenuation of collagen degradation as part of a path to stabilize lung function.

## Methods

### Sex as a biological variable

Sex was not considered as a biological variable. Lung donors of both sexes were obtained in the study.

### Human Lung Tissue and bronchoalveolar lavage (BAL)

Tissues from normal lungs declined for transplantation and from IPF patients undergoing transplantation were deidentified and donated to research through institutional protocols approved by the UCSF Institutional Review Board.

BAL fluids were generated by instilling 100 ml of PBS into a whole lobe. BAL pellets were produced by centrifugation of BAL fluids at 500g for 10 minutes and then lysed in RIPA buffer for Western blotting. The list of antibodies is provided in Supplemental Table 4.

### Human lung protein isolation and mass spectrophotometry

Human lung tissue was homogenized in RIPA buffer (ThermoFisher 89900) with protease and phosphatase inhibitors (ThermoFisher 78440) as previously described (35). For mass spectrophotometry, whole lung lysate was incubated with HTII-280 hybridoma supernatant (Terrace Biotech) or anti-mouse IgM antibody as a control (Abcam) then 100ul of Protein L magnetic beads (Pierce 88849) were added to the mixture. After several washes using TBST+300uM NaCl, the beads were incubated in immunoblot loading buffer (Biolegend cat#. 426311). The resulting eluates (HTII-280 and IgM) were used in mass spectrometry analysis. *Analysis of Mass Spectrometry data.* The immunoprecipitated HTII-280 and IgM control samples were run on a stain free gel and isolated areas of the gel corresponding to the hypothesized molecular weight, 280 kD. After running mass spectrometry on these samples, both spectral counts and area under curve for peptides in the IgM and HTII-280 samples were counted. After results were obtained, all proteins with >2 unique peptides recognized and >2-fold enrichment in area + spectral counts in HTII-280 sample/IgM only were reported.

### Immunoprecipitation and Western Blot

Pulverized PCLS tissues lysed in RIPA buffer and HTII-280 immunoprecipitate were analyzed by immunoblotting as previously described (35). For MUC1 immunoprecipitation, 1mg of whole lung lysate was first incubated with 4ug MUC1 antibody (Santa Cruz) or 4ug anti-mouse IgG antibody (ThermoFisher), then 50ul of Protein A/G magnetic beads (ThermoFisher cat # 80104G) were added to the mixture. After several washed in TBST+300uM NaCl, the beads were incubated in immunoblot loading dye (Biolegend cat# 426311). The resulting eluate was directly used for immunoblotting with HTII-280 (1:50).

Densitometry was quantified using NIH ImageJ software. The list of antibodies is provided in Supplemental Table 4. Samples were loaded onto a gel based on equalizing all samples among their total protein concentrations and analyzed by immunoblotting. PCLS conditioned medium (CM) was concentrated ten times and protein concentration was measured. Densitometry was quantified using NIH ImageJ software. BAL cell lysate Western blot quantification was performed by normalizing band intensity of each sample to b-actin of that sample. 10X PCLS-CM Western blot quantification was performed by normalizing band intensity of each sample to total protein in CM of that sample.

### Deglycosylation of human lung tissue

Human lung tissue lysates from three different donors were treated with neuraminidase, PNGaseF, and O-deglycosidases according to the instructions in the NEB Protein Deglycosylation Mix II protocol. Resulting deglycosylated lysate was added to 5x loading buffer with BME and used for immunoblotting with HTII-280.

### Immunofluorescence and quantification: 1) Optimal Cutting Temperature (OCT) embedding

Lungs and organoids were fixed with 4% PFA (cat. no. 15714 Electron Microscopy Sciences), and embedded in OCT after a sucrose gradient, as previously described (21). Sections (7μm) were cut on a cryostat. 2) *Cytospins of sorted cells*. Cell pellets were fixed in 4%PFA then loaded into chambers and spun onto superfrost plus microscope slides (cat. no. 12-550-15, Thermo Fisher). Slides were subsequently immunostained. 3) *Immunofluorescent staining.* Prior to antigen retrieval, OCT-embedded slides were fixed in 4% PFA. Antigen retrieval (cat. no.

DV2004MX, Biocare) was performed. Slides were washed with PBS, blocked/permeabilized (5% horse serum 0.5% BSA 0.1% Triton X) for 1 hour, and then incubated with primary antibodies overnight at 4°C and with Alexa Fluor secondary antibodies for 1 hour. (Supplemental Table 4). Images were captured using ZEN v3.1 software (Zeiss); 3) *Image quantification.* Slides were imaged for quantification on a Zeiss AxioImager.M1 microscope. Cell counts for stained organoids or cytospins were performed manually. Approximatively 50 organoids or 1,000 cells (cytospin) per condition were counted blindly by members of the laboratory. The results were averaged between each specimen and s.d. values were calculated per condition.

### Lung tissue processing and fluorescence activated cell sorting. 1) Lung digestion, fluorescence activated cell sorting (FACS)

A single cell preparation of normal, ILD explant, or biopsy tissues was prepared as previously described (10). FACS was performed on digested donor lung tissue to obtain AT2 cells (EPCAM^+^/CD11b^neg^/CD31^neg^/ CD45^neg^/ HTII-280^+)^, mesenchymal cells (EPCAM ^neg^/CD11b^neg^/CD31^neg^/CD45 ^neg^), basal cells (EPCAM^+^/CD11b^neg^/CD31^neg^/CD45 ^neg^/ HTII-280^neg^/NGFR^+^), and ceacam6 cells (EPCAM^+^/CD11b^neg^/CD31^neg^/CD45 ^neg^/ HTII-280^neg^/CD66c^+^/NGFR^neg^).

### In vitro and ex vitro models. 1) Cell culture

Adult human mesenchymal cells were cultured in DMEM (Cat#11965092, Thermo Fisher) with 10% fetal bovine serum (Cat#SH883IH2540, Fisher Scientific), 1% GlutaMAX (Cat#35050-61, Gibco), 1% HEPES (Cat#5630-080, Gibco), and 1% Pen/Strep (Cat#10378016, Gibco). Cells were used within the first five passages of being isolated from donor lungs for mesenchymal cells. The 293 cells (Cat#CRL1573, ATCC) were cultured in DMEM supplemented with penicillin/streptomycin, and 10% FBS. All the cell lines in the laboratory are periodically tested for mycoplasma contamination. Only the mycoplasma-free cells are used for experiments. Basal cells were infected with an mCherry lentivirus (UCSF viral core) at an MOI of 10 with LentiTrans (cat. no. LTDR1) and 10uM y-27632 compound (cat. no. 1254) overnight and then expanded in PneumaCult™-Ex Plus Medium (cat# 05040, StemCell Technologies). 2) *Organoid assay*. AT2s or CD66c+/NGFR ^neg^ or NGFR+ cells and mesenchymal cells were co-cultured (5,000 AEC2s or CD66c^+^ cells: 30,000 fibroblasts/mesenchymal cells per well) in modified MTEC medium diluted 1:1 in growth factor-reduced Matrigel (cat# CB-40230A, Thermo Fisher). Modified MTEC culture medium is composed of SABM with insulin, transferrin, bovine pituitary extract, retinoic acid and epidermal growth factor (EGF) as per the SAGM Bullet Kit and 0.1 μg.ml^−1^ cholera toxin (cat# C8052, Sigma), 5% charcoal treated FBS and 1% Pen/Strep. The cell suspension–Matrigel mixture was placed in a transwell and incubated with 10 μM y-27632 compound (cat# 1254, Tocris) for the first 24 hours. Isolation of the cells from organoids was performed as previously described (17). Each experimental condition was performed at least in triplicate or as indicated in figure legends. Where applicable, Frizzled-5 agonist (Fzd5Ag, 25nM, AntlerA, Toronto, Ontario Canada) was added to the medium and replenished in every medium change for the blocking (day 1 to day 14) experiments and added to the medium on day 11 until day 19 for the reversal experiments. Organoids were processed for OCT-embedding for immunostaining or made into a single cell preparation and positively selected for EPCAM for RNA extraction. When the AT2-derived basal cells were used for xenotranplant, the cells were freshly flow sorted for EPCAM prior to transplantation.

### Spatial transcriptomic (Xenium)

Human lung tissue sections were obtained from donor lungs and explanted lungs of patients with usual interstitial pneumonia (UIP) associated with idiopathic pulmonary fibrosis (IPF), scleroderma, rheumatoid arthritis (RA), or hypersensitivity pneumonitis (HP). Tissue sections were processed using the Xenium platform (10x Genomics) and hybridized with a selected panel of RNA probes for *COL1A1*, *MMP1*, *ABCA3*, *SCGB3A2*, *KRT5*, and *KRT1*7. Cell segmentation was performed with Xenium in situ multimodal cell segmentation. The Xenium output files were loaded into R and analyzed using the Seurat package. Data were log-normalized and analyzed following the Seurat spatial data vignette (https://satijalab.org/seurat/articles/seurat5_spatial_vignette_2). Cell annotation was performed using a custom R script that integrated gene expression levels to assign cell identities. Quantification of cell populations within respiratory bronchioles and microcystic regions was conducted by manually selecting anatomical zones using Xenium Explorer software. Annotated cells were then counted within these regions using the integrated quantification tool of the Xenium Explorer software.

*Single-cell RNA analysis:* was performed as previously described (17).

### RNA extraction and Quantitative RT-PCR

RNA was extracted using the ReliaPrep RNA Cell Miniprep System (cat# Z6011, Promega) as per manufacturer’s instructions. Reverse transcription was performed with iScript RT Supermix (cat# 1708841 Bio-Rad) and quantitative real-time PCR (qPCR) was performed using SsoAdvanced Univ SYBR Green Suprmix (cat# 1725271 Biorad). Relative expression was calculated with the delta-delta method. The list of primers is provided in Supplemental Table 5.

### Collagen Degradation Assay

DQ™ bovine skin type I collagen (fluorescein-conjugated) was purchased from <u>ThermoFisher</u>. Conditioned medium from 8-day basal cell culture or 5-day PCLS culture were incubated with DQ-collagen I (50 ug.ml^-1^) for 1 hour at 37°C in a black 96-well plate. MMP activity was determined by fluorescence intensity measured using a microplate reader (ex/em = 495/515 nm).

### Xenotransplantation Assay

300,000 to 500,000 AT2-derived basal cells or mCherry labeled IPF basal cells in 40 µL volume (1XPBS) were transplanted at 10 days post bleomycin injury into the lungs of NOD *scid* gamma mice via oral aspiration. A subgroup of mice was treated with the pan MMP inhibitor GM6001 (10mg.kg^-1^, cat# HY-15893, MedChemExpress) the same day as the transplant until the end of the protocol. Transplanted mice were euthanized 10 days post-transplant (a total of 20 days post-bleomycin injury). After euthanasia, 2 x 1 ml of PBS (BALF) were injected into the lungs, collected, centrifuged at 500g for 10 minutes and then lysed in RIPA buffer for Western blotting. *Precision Cut Lung Slices.* Fresh lung tissues were obtained from xenograft transplanted mice. Precision-cut lung slices (PCLS) were prepared as described (12). Lung tissues were inflated with warm 2% low-melting agarose (cat# 16550100, ThermoFisher) and placed in cold PBS. Solidified tissues were cut into 400µm thick slices using Compresstome (cat# VF-310-0Z; Precisionary Instrument LLC.) Ten randomly selected slices per well of a 6-well plate were cultured in serum-free DMEM (Dulbecco’s modified Eagle’s medium) supplemented with 100 units.ml^-1^ penicillin and streptomycin under standard cell culture conditions (37°C, 5% CO2, 100% humidity) with Nystatin (cat# N186, ThermoFisher, 20U.ml^-1^. At day 5, all cultured PCLSs were immediately transferred into liquid nitrogen and subsequently stored at −80 °C prior to protein extraction.

Animal experiments were conducted using a protocol approved by the University of California, San Francisco, Institutional Animal Care and Use Committee (Protocol AN109566).

### NFATc3 nuclei translocation assay

The 293 cells with 80% confluency were co-transfected with FZD5 plasmid (GenScript) using TurboFect transfection reagent (cat# R0531, Thermo Fisher) and cultured in DMEM complete medium for 24-48 hours. Cells were cultured serum-free medium with or without SFRP2 (30 ng.ml^−1^) and with or without Fzd5Ag (100 nM) for 1 hour at 37 °C before lysis for nuclei isolation. After cell lysis, nuclei pellet was washed with lysis buffer once before solubilizing in RIPA buffer. Clarified nuclei supernatants along with cytosol control were blotted for NFAT3, FZD5, GAPDH, or NUP62 (Supplemental Table 4).

### Statistics

Statistical analyses for cell count and gene and protein expression were performed in GraphPad Prism. The two-tailed Mann-Whitney test was used for statistical analysis of scRNA-Seq gene signature expression. One-Way ANOVA, unpaired and paired two-tailed t-tests were used to determine the P values, and the data in the graphs are presented as mean ± s.d.

Unpaired t-test was used to compare two treatment groups. The Kruskal–Wallis test or Dunnett’s multiple comparisons test were used for multiple comparisons. For normally distributed data, ordinary one-way ANOVA followed by Tukey’s multiple comparisons test was performed.

### Data Availability

Raw data files are available on the Mass Spectrometry Interactive Virtual Environment, a member of the Proteome Xchange consortium, under the identifier MSV000099867. Previously published scRNA-seq data that are re-analysed here are available at <u>GSE150068</u> and <u>GSE150247</u>. Publicly available R packages were used for all computational analyses. No custom codes were developed. Representative code is available on reasonable request. Data used for graphing/quantification purposes are provided in the Supporting Data Values file. All raw data used for quantification and statistical analysis are available in a single GraphPad Prism file that can be made available upon request.

## Author Contributions

AJ, CJLS, and HAC conceived elements of the study. KB, ANB, MT, AJ, YW, TCH, VR, BZ, SI, and CJLS performed the experiments. AJ, VR, BZ, and DS conceived and executed all the experiments related to HTII-280/MUC1. MM and PJW provided human lung samples. KG, ANB, MT, AJ, CJLS, and HAC wrote the manuscript. All authors read and reviewed the manuscript.

KG and ANB were designated co-first authors based on their relative overall contributions to performing and analyzing the cell biology aspects of the study. KG was listed first for her initial bioinformatic analyses underlying the conception of this study.

## Funding support

This work is supported by NIH grants R35HL150767 and U01HL134766 and California Institute for Regenerative Medicine grant DISC0-13788 (H.A.C.), NIH grants T32HL007185 (AJ, SI), F32HL172536 (AJ), F32HL175915 (SI), K08HL157654 (VA), American Lung Association grant HIA-1268508 (VA) and by the Nina Ireland Program Award for human lung collection (M.M.).

Biorender was used in figure preparation.

## Acknowledgements

The authors thank Dr. Erin Gordon for sharing cultures of airway basal cells and Mazharul Maishan for his assistance in the access of normal lung donors.

## Supplemental material

**1-Supplemental methods**

**2-Supplemental Tables**

**3-Supplemental Figures**

## Supplemental Methods

*Human lung protein isolation and Immunoprecipitation:* Human lung tissue was homogenized in RIPA buffer (ThermoFisher 89900) with protease and phosphatase inhibitors (ThermoFisher 78440). For mass spectrophotometry, 1mg of whole lung lysate was first incubated with 2ug HTII-280 hybridoma supernatant (Terrace Biotech) or 2ug anti-mouse IgM antibody as a control (Abcam) for 45 minutes, then 100ul of Protein L magnetic beads (Pierce 88849) were added to the mixture and incubated for an additional 45 minutes. A magnetic stand (Dynal MPC-S) was used to bind the beads for 5 minutes, and the beads were washed 7 times with wash buffer (TBST+300uM NaCl). Afterwards, the beads were incubated in immunoblot loading buffer (Biolegend cat#. 426311). The resulting eluates (HTII-280 and IgM) were used in mass spectrometry analysis. *Analysis of Mass Spectrometry data.* The immunoprecipitated HTII-280 and IgM control samples were run on a stain free gel and isolated areas of the gel corresponding to the hypothesized molecular weight, 280 kD. After running mass spectrometry on these samples, both spectral counts and area under curve for peptides in the IgM and HTII-280 samples were counted. After results were obtained, all proteins with >2 unique peptides recognized and >2-fold enrichment in area + spectral counts in HTII-280 sample/IgM only were reported.

*In-gel Digestions for Mass Spectrometry Analysis*: Immunoprecipitation (IP) samples were diluted in 4x Laemmli Sample Buffer (BioRad) containing 10% β-mercaptoethanol. Samples were loaded onto a protein gel (BioRad, 4-15% Criterion TGX Stain-Free Precast Gel) and proteins were separated at 200V for 42 minutes. Gels were stained with colloidal blue stain (Invitrogen) overnight at room temperature with gentle agitation. Excess stain was removed by briefly rinsing the gel in de-stain solution (50% water, 40% methanol, 10% acetic acid) for 3 minutes followed by 3 x 15 min washes in pure water. Each lane was excised (56-280 kDa) out of the gel and then each lane was further segmented using fresh razor blades. Each gel slice was subsequently diced into approximately 1 mm cubes, and this was then transferred to a 1.5 ml microcentrifuge tube (Protein Lo-Bind, Eppendorf). 100 mM Ammonium Bicarbonate (ABC) was added to each tube (75 μl, or enough to completely cover gel cubes), and samples were left incubating at room temperature for 10 minutes. ABC solution was removed with gel-loading tips (VWR). Then, 200 μL of ABC (100 mM) and 20 μL of DTT (50 mM) were added to each sample before incubating at 55°C (500 rpm, for 30 minutes). Excess buffer was removed with a gel-loading tip and replaced with 200 μL of ABC (100 mM) and 20 μl of iodoacetamide (150 mM). Samples were incubated at room temperature in the dark for 30 minutes. Buffer was removed, and gel cubes were washed 4x with 150-300 μL of 1:1 ABC (100 mM):Acetonitrile on a rotator at room temperature for 10 min. After the last wash, gel cubes were dehydrated with 150 μl of acetonitrile. Excess acetonitrile was removed on a speed-vac for 50 minutes. A cocktail of trypsin (Promega) at 3.3 ng.μl^-1^ in ABC (50 mM) was made and 150 μl of this was added to each sample (0.5 ug trypsin/sample). Samples were left digesting at 37°C at 500 rpm for 16 hours.

After digestion, samples were acidified and peptides eluted through the addition of 250 μl of a 66% acetonitrile, 33% ABC (100 mM), 1% formic acid solution. Samples were then centrifuged at 10,000 rcf for 2 min, and the resulting supernatants were transferred to new Protein LoBind eppi tubes. Peptide elution step was repeated 1x and corresponding supernatants were combined. Samples were then dried on a speed-vac for 2 hours. Crude dried peptides were then de-salted using ZipTip C18 columns (Millipore) and then dried down once more on a speed-vac for 20 minutes. The resulting peptides were then re-suspended in loading buffer (2% ACN, 0.1% FA in water) and completely solubilized in a bath sonicator prior to mass spectrometry analysis.

*Mass spectrometry data acquisition and analysis:* A nanoElute was attached in line to a timsTOF Pro equipped with a CaptiveSpray Source (Bruker, Hamburg, Germany). Chromatography was conducted at 40°C through a 25cm reversed-phase C18 column (PepSep) at a constant flow rate of 0.5 μl.min^-1^. Mobile phase A was 98/2/0.1% Water/ACN/Formic Acid (v/v/v) and phase B was ACN with 0.1% Formic Acid (v/v). During a 108 min method, peptides were separated by a 3-step linear gradient (5% to 30% B over 90 min, 30% to 35% B over 10 minutes, 35% to 95% B over 4 minutes) followed by a 4 minutes isocratic flush at 95% for 4 minutes before washing and a return to low organic conditions. Experiments were run as data-dependent acquisitions with ion mobility activated in PASEF mode. MS and MS/MS spectra were collected with *m*/*z 1*00 to 1700 and ions with *z* = +1 were excluded.

Raw data files were searched using PEAKS Online Xpro 1.6 (Bioinformatics Solutions Inc.). The precursor mass error tolerance and fragment mass error tolerance were set to 20 PPM and 0.05 respectively. The trypsin digest mode was set to semi-specific and missed cleavages was set to 2. The human Swiss-Prot reviewed (canonical) database (downloaded from UniProt) and the common repository of adventitious proteins (cRAP, downloaded from The Global Proteome Machine Organization) totaling 20,487 entries were used. Carbamidomethylation was selected as a fixed modification. Acetylation (N-term), Deamidation (NQ) and Oxidation (M) were selected as variable modifications. The maximum number of variable PTMs per peptide was 3.

### Immunoprecipitation

For MUC1 immunoprecipitation, 1mg of whole lung lysate was first incubated with 4ug MUC1 antibody (Santa Cruz) or 4ug anti-mouse IgG antibody (ThermoFisher) for 45 minutes, then 50ul of Protein A/G magnetic beads (ThermoFisher cat # 80104G) were added to the mixture and incubated for an additional 45 minutes. A magnetic stand (Dynal MPC-S) was used to bind the beads for 5 minutes, and the beads were washed 7 times with wash buffer (TBST+300uM NaCl). Afterwards, the beads were incubated in immunoblot loading dye (Biolegend cat# 426311). The resulting eluate was directly used for immunoblotting with HTII-280 (1:50). The list of antibodies is provided in Supplemental Table 4.

### Immunofluorescence and quantification: 1) Optimal Cutting Temperature (OCT) embedding

Lungs inflated with 94%OCT/2%PFA/4%PBS and organoids in 3D Matrigel were fixed with 4% PFA (cat. no. 15714 Electron Microscopy Sciences), washed with PBS three times and embedded in OCT after 30% sucrose and 15% sucrose/50% OCT gradient washing (overnight for lungs, 30 minutes for organoids). Sections (7-μm) were cut on a cryostat. 2) *Cytospins of sorted cells*. Cell pellets were resuspended in 50ul per chamber in 4%PFA and incubated for 10 minutes at room temperature with intermittent vortexing. Cells were loaded into chambers and spun onto superfrost plus microscope slides (cat. no. 12-550-15, Thermo Fisher) using a StatSpin Cytofuge 2 cytocentrifuge. Slides were washed 2x in PBS and subsequently immunostained. 3) *Immunofluorescent staining.* Prior to antigen retrieval, OCT-embedded slides were fixed in 4% PFA for 10 minutes, then washed with PBS and cytospins were washed with PBS. Antigen retrieval (cat. no. DV2004MX, Biocare) was performed for 20 minutes in an oven at 155°C and washed in distilled water. Slides were washed with PBS, blocked/permeabilized (5% horse serum 0.5% BSA 0.1% Triton X) for 1 hour, and then incubated with primary antibodies overnight at 4°C (Supplemental Table 4). Slides were washed with PBS and then incubated with Alexa Fluor secondary antibodies for 1 hour. 4′,6-Prior to mounting, diamidino-2-phenylindole (DAPI) was added for 5 minutes and slides were mounted with prolong gold.

Images were captured using ZEN v3.1 software (Zeiss); 3) *Image quantification.* Slides were imaged for quantification on a Zeiss AxioImager.M1 microscope. Cell counts for stained organoids or cytospins were performed manually. Approximatively 50 organoids or 1,000 cells (cytospin) per condition were counted blindly by members of the laboratory. The results were averaged between each specimen and s.d. values were calculated per condition.

### Lung tissue processing and fluorescence activated cell sorting. 1) Lung digestion, fluorescence activated cell sorting (FACS)

A single cell preparation of normal, ILD explant, or biopsy tissues was made using mechanical disruption and enzymatic digestion (dispase,15 I.U.ml^-1^ and collagenase, 225u.ml^-1^).

### Single cell preparation from organoids

The cell–Matrigel mixture in the transwell was washed with PBS and incubated in the 15u.ml^-1^ dispase for 30 minutes at 37°C with intermittent resuspension. The mixture was removed from the transwell and resuspended in TrypLE (cat# 12563011, ThermoFisher). Cells were shaken at 37 °C for up to 20 minutes, pipetting up and down 10 times every 5 minutes and checking for single cells. For the organoids isolation only, cells were then stained with biotin anti-CD326 (cat# 324216, BioLegend) for 30 minutes at 4°C. Streptavidin beads (cat# 17663, STEMCELL) were added to isolate the epithelial cells, and the rest of the cells were mesenchymal cells. AT2-2D cells were washed twice with PBS. Dispase (15U.ml^-1^) was added, and plate was incubated for 35 minutes shaking at 37°C. Dispase was carefully collected from the wells without disturbing the matrigel. Wells were washed twice with PBS to ensure recovery of all cells.

### NFATc3 nuclei translocation assay

The 293 cells with 80% confluency were co-transfected with FZD5 plasmid (GenScript) using TurboFect transfection reagent (cat# R0531, Thermo Fisher) and cultured in DMEM complete medium for 24-48 hours. Cells were washed with PBS 3 times and replaced with serum-free medium with or without SFRP2 (30 ng.ml^−1^) and with or without Fzd5Ag (100 nM) for 1 hour at 37 °C before lysis for nuclei isolation. The 293 cells were lysed in ice-cold NP40 lysis buffer (5 mM Tris, pH 8.0, 15 mM NaCl, and 0.1% NP40) supplemented with protease/phosphotase inhibitors and 1 mM phenylmethylsulfonyl fluoride. After 5 minutes incubation with lysis buffer, cells were scraped off and centrifuged at 500 g for 5 minutes at 4°C. Supernatant was saved as cytosol control. Nuclei pellet was washed with lysis buffer once before solubilizing in RIPA buffer. Clarified nuclei supernatant along with cytosol control were blotted for NFAT3, FZD5, GAPDH, or NUP62 (Supplemental Table 4).

*Single-cell RNA analysis*. FASTQ files were run through CellRanger v2.1.1 software with default settings for de-multiplexing, aligning reads with STAR software to Hg19 or GRCh38, and counting unique molecular identifiers (UMIs). Seurat package v5.3.0 in R v4.3.1 was used for downstream analysis. Low-quality cells were filtered (expressing fewer than 200 genes, >10% mitochondrial reads and >6,000 unique gene counts). Principal component analysis was performed on log-normalized and integrated data using 2,000 variable genes. The top 10 principal component analyses were used for clustering and visualized using the UMAP algorithm in the Seurat package. Pathway activity analysis was done using the decoupleR package. Single-cell transcriptomes were obtained from <u>GSE150068</u> (organoids and IPF) and <u>GSE150247</u> (normal human lung) and processed using Seurat. Basal cells were extracted and analyzed. The lists of DEGs were identified with a non-parametric Wilcoxon rank sum test.

---

**Supplemental Table 1:**
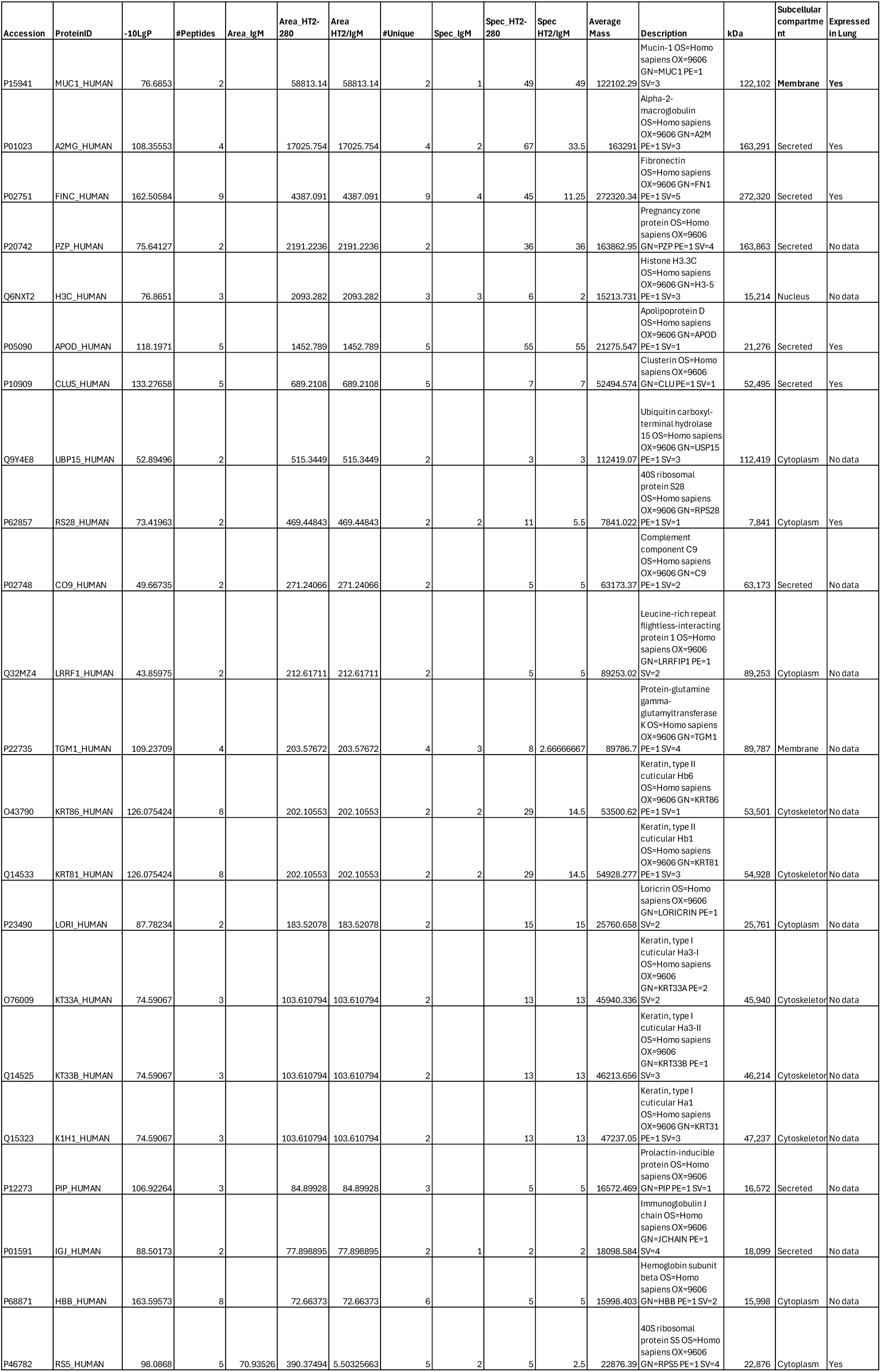
List of top proteins co-IP with HTII-280 identified by mass spectrophotometer.

**Supplemental Table 2:** Characteristics of patients included in spatial transcriptomic analysis (Xenium).

| **Patients** | **Age** | **Sex** | **Diagnosis** | **Pathology** |
| --- | --- | --- | --- | --- |
| 1 | 31 | M | Control | Normal |
| 2 | 78 | F | Control | Normal |
| 2 | 48 | M | RA | UIP and NSIP |
| 3 | 59 | F | HP | UIP |
| 4 | 68 | F | IPF | UIP |
| 5 | 65 | M | IPF | UIP |
| 6 | 64 | M | IPF | UIP |

**Supplemental Table 3:** Single-cell differential gene expression of basal cells from control and IPF lungs (n = 2 normal and 4 IPF).

| Gene | avg_log2FC | pct.1 | pct.2 | p_val_adj |
| --- | --- | --- | --- | --- |
| *ACKR1* | 9.3770164 | 0.184 | 0.001 | 1.60E-111 |
| *AC131056.5* | 9.3597975 | 0.145 | 0 | 1.38E-86 |
| *HBA2* | 8.8117018 | 0.128 | 0.005 | 9.01E-68 |
| *HBB* | 8.1243594 | 0.135 | 0.007 | 9.14E-69 |
| *MIF-AS1* | 7.6456736 | 0.218 | 0.007 | 1.13E-125 |
| *RP11-138I1.4* | 6.1642181 | 0.102 | 0.006 | 1.86E-49 |
| *MMP1* | 5.9744862 | 0.621 | 0.086 | 0.00E+00 |
| *AC090498.1* | 5.3457207 | 0.978 | 0.53 | 0.00E+00 |
| *UQCRHL* | 5.0543669 | 0.345 | 0.053 | 7.87E-161 |
| *RP5-940J5.9* | 4.9402 | 0.171 | 0.017 | 7.22E-78 |
| *LGALS7* | 4.7993151 | 0.215 | 0.063 | 6.30E-60 |
| *BEST1* | 4.7299151 | 0.331 | 0.044 | 3.80E-159 |
| *CTC-250I14.6* | 4.6258266 | 0.113 | 0.016 | 1.86E-42 |
| *TNFSF9* | 4.4815466 | 0.103 | 0.015 | 1.53E-37 |
| *SNX22* | 4.4051164 | 0.122 | 0.016 | 2.28E-48 |
| *HSPA6* | 4.3759943 | 0.168 | 0.032 | 2.65E-57 |
| *IFITM1* | 4.3705056 | 0.778 | 0.219 | 0.00E+00 |
| *NME2* | 4.2486114 | 0.539 | 0.138 | 4.45E-255 |
| *PAM16* | 4.1257975 | 0.107 | 0.02 | 7.56E-35 |
| *RP11-386G11.10* | 4.1214759 | 0.123 | 0.02 | 1.99E-43 |
| *TPPP3* | 4.0366815 | 0.564 | 0.156 | 6.54E-259 |
| *ZNF90* | 3.9936646 | 0.628 | 0.231 | 1.67E-276 |
| *LGALS7B* | 3.8537267 | 0.634 | 0.289 | 6.89E-252 |
| *HNRNPA1L2* | 3.8163356 | 0.44 | 0.116 | 1.28E-176 |
| *NACA2* | 3.8020716 | 0.34 | 0.101 | 8.24E-111 |

**Supplemental Table 4:**
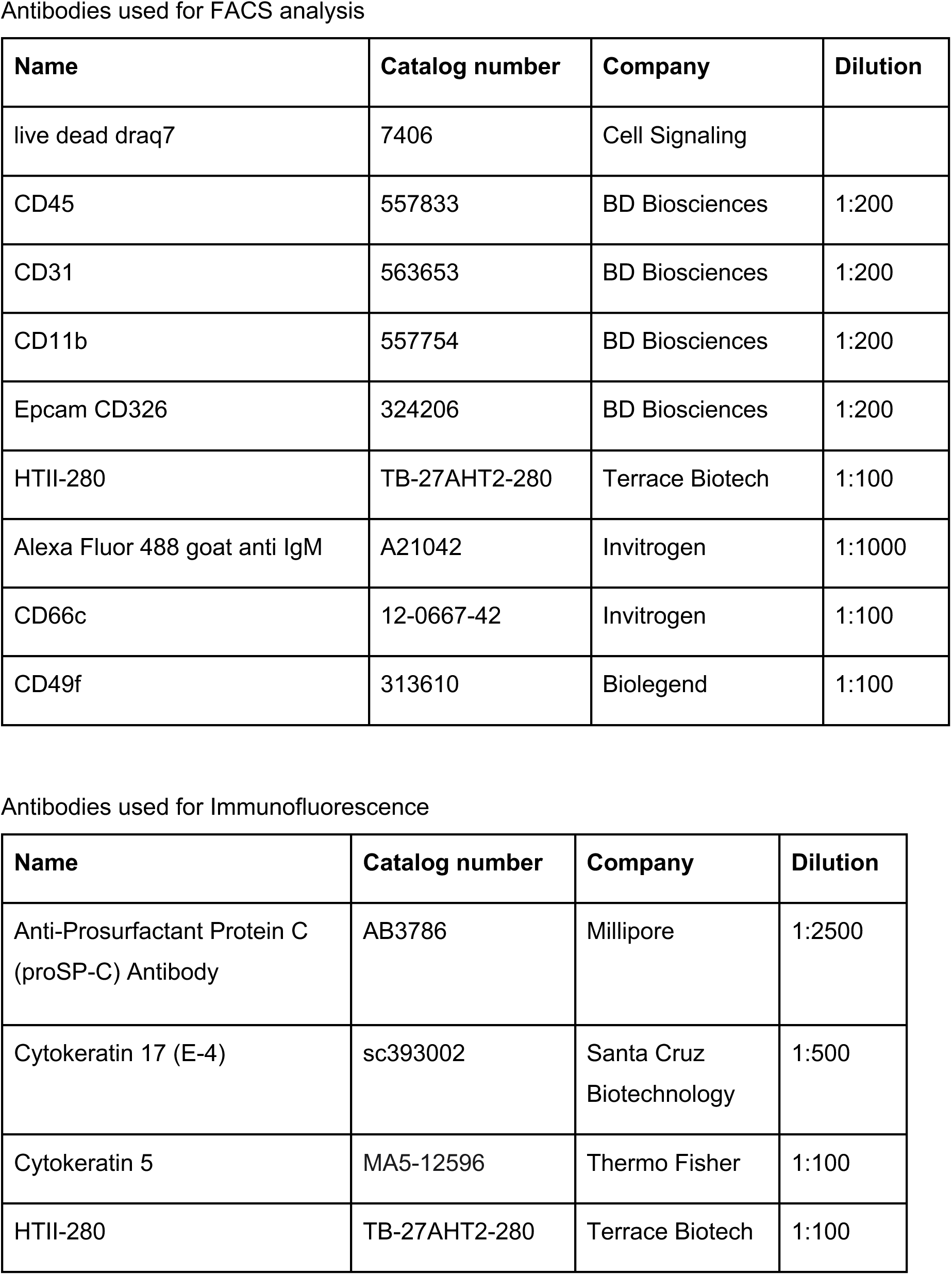
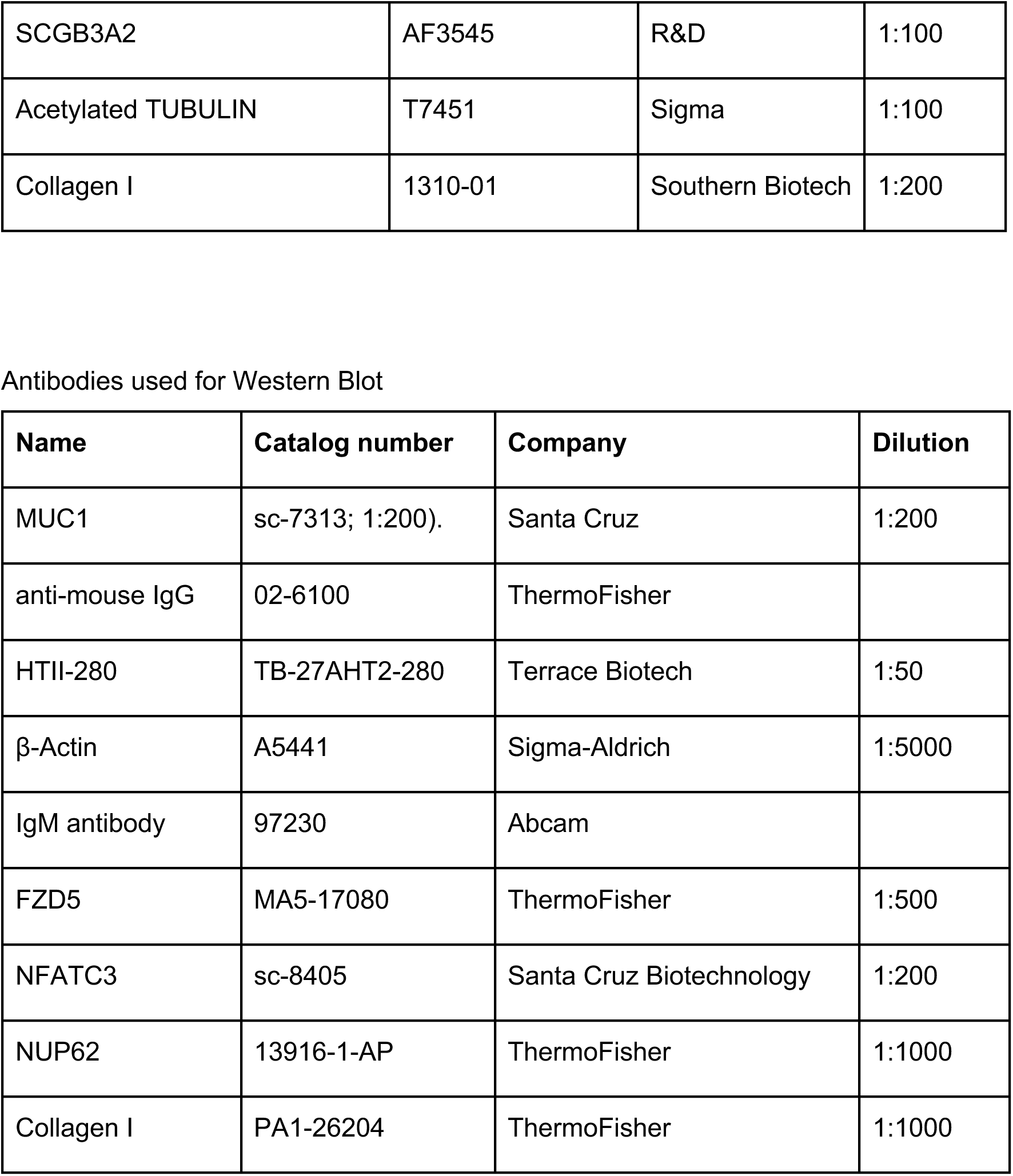
List of antibodies used for cell sorting FACS, western blot, and immunofluorescence.

**Supplemental Table 5:**
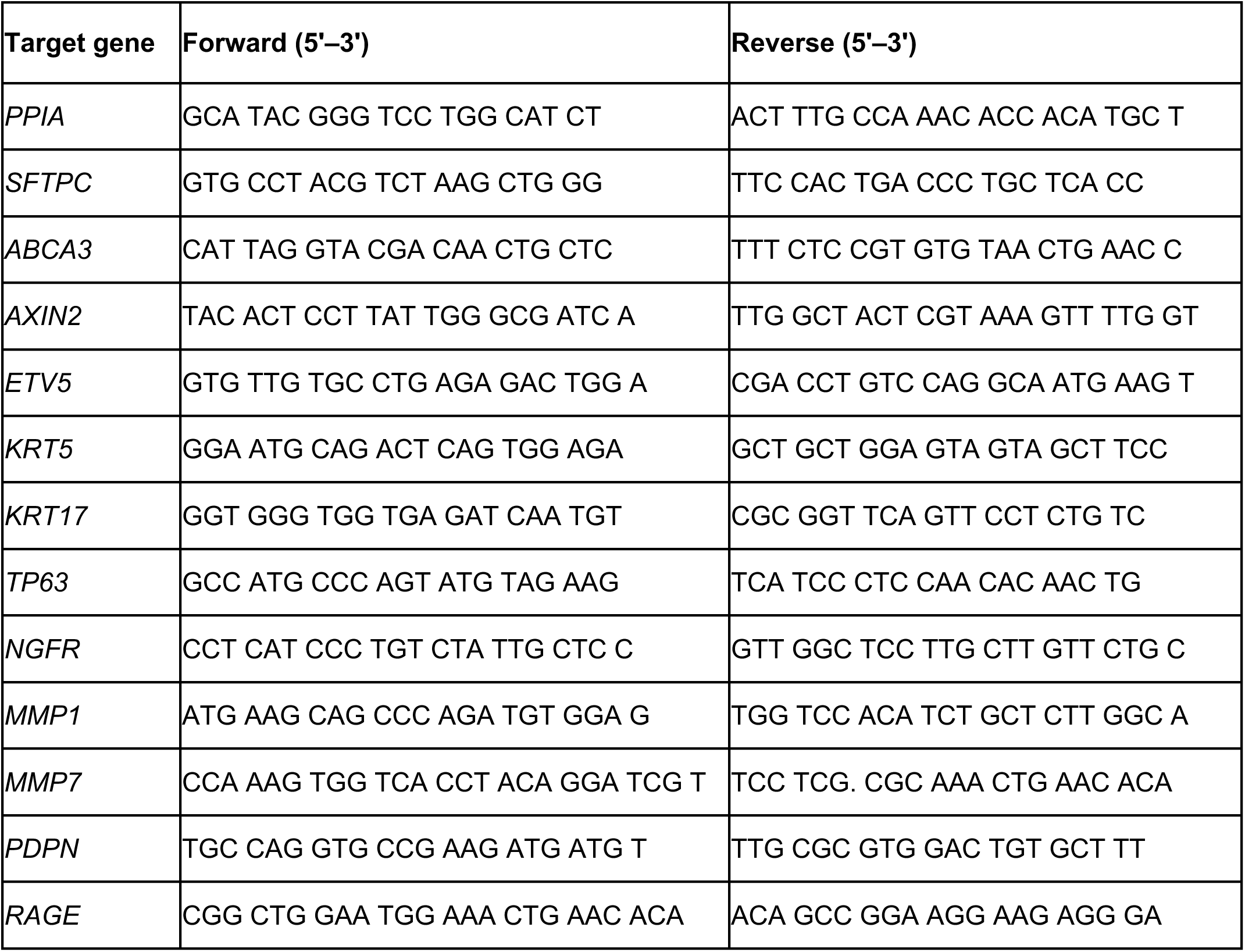
List of human primers used for qPCR.

**Supplemental Figure 1:**
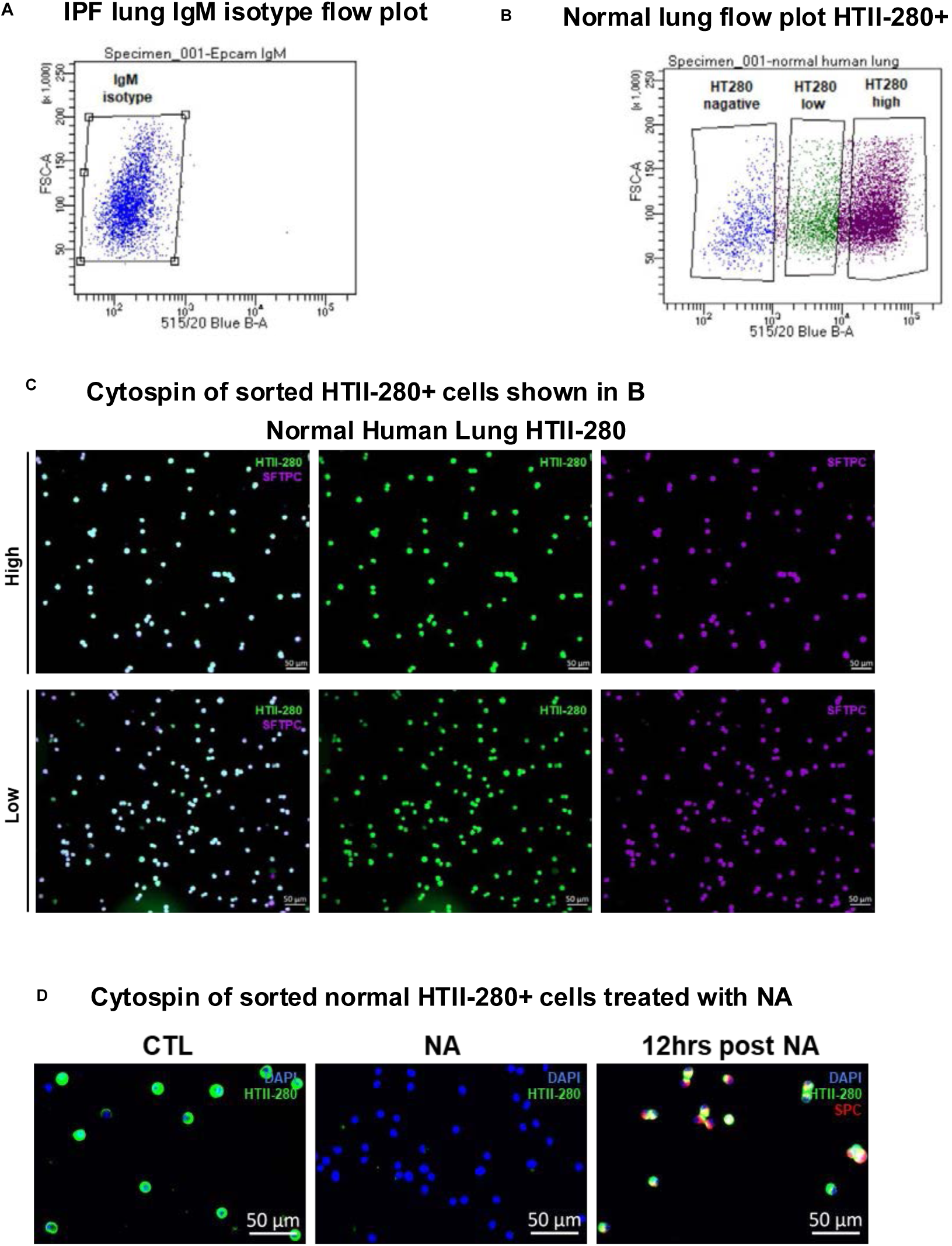
(**A**) Representative flow blot of epithelial cells stained with the IgM isotype control from an IPF lung. (**B**) Representative flow blot of sorted HTII-280^+^ cells indicative of two distinctive populations, HTII-280^low^ and HTII-280^hi^ cells from a normal lung above the MFI of the isotype control. (**C**) Representative images of HTII-280^hi^ and HTII-280^low^ cytospins from a normal lung, stained by IF for HTII-280 (green), or SFTPC (purple). (**D**) Representative images of HTII-280 cytospins ± NA from a normal lung, stained by IF for HTII-280 (green), or SFTPC (red).

**Supplemental Figure 2:**
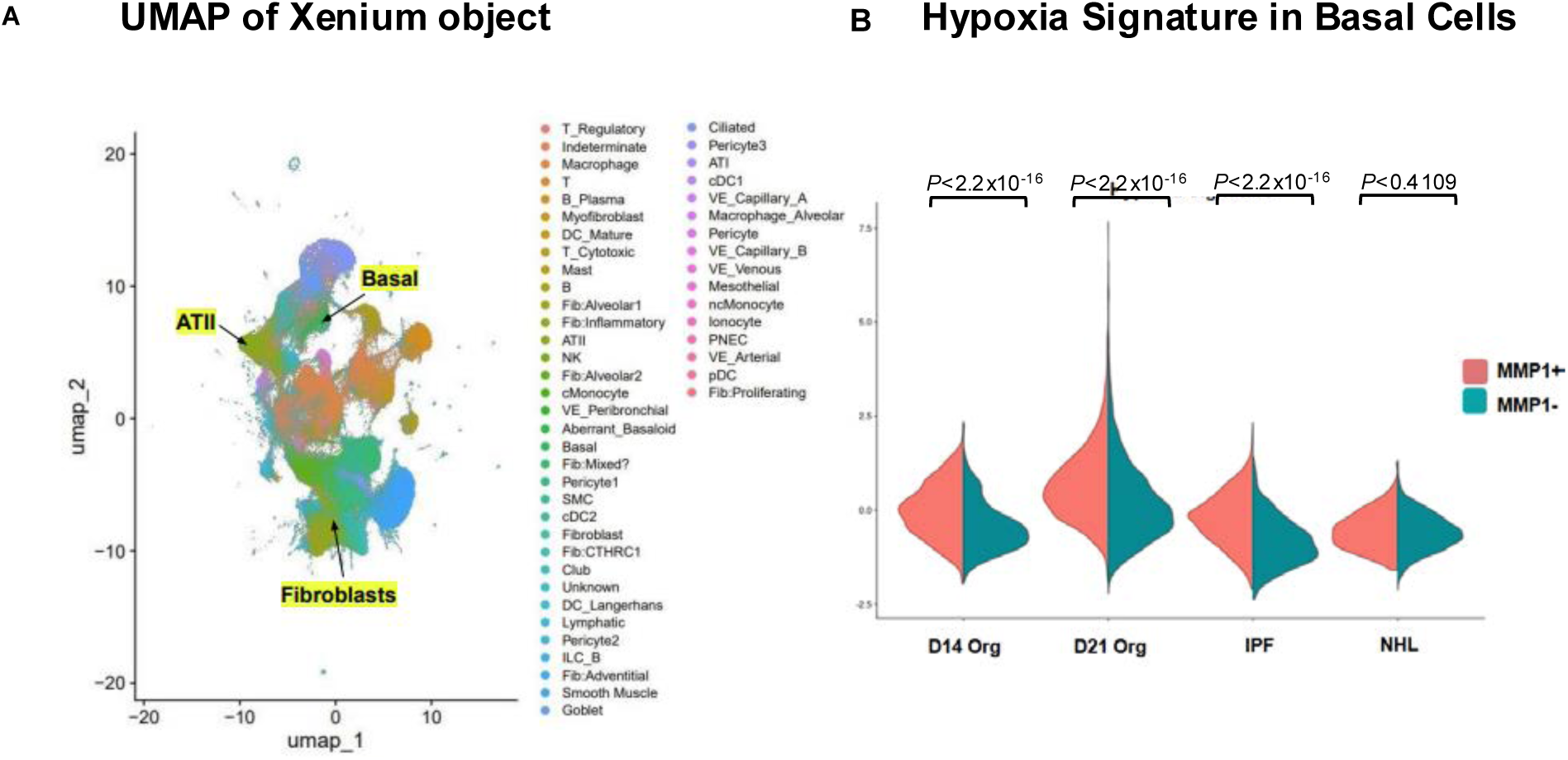
(**A**) UMAP grouped by cell types of single-cell RNA sequencing control and IPF human lung from spatial transcriptomic probe set (n = 7 IPF and 3 control biological replicates). (**B**) Violin plot of Hypoxia signature in MMP1^neg^ or MMP1^+^ basal cells from normal, IPF lungs, and from organoids after 14 and 21 days (n =2 normal lung, 4 IPF lungs, 1 set of organoids collected at day 14, and 1 set of organoids collected at day 21). Statistical analysis was determined by Mann-Whitney t-test.

**Supplemental Figure 3:**
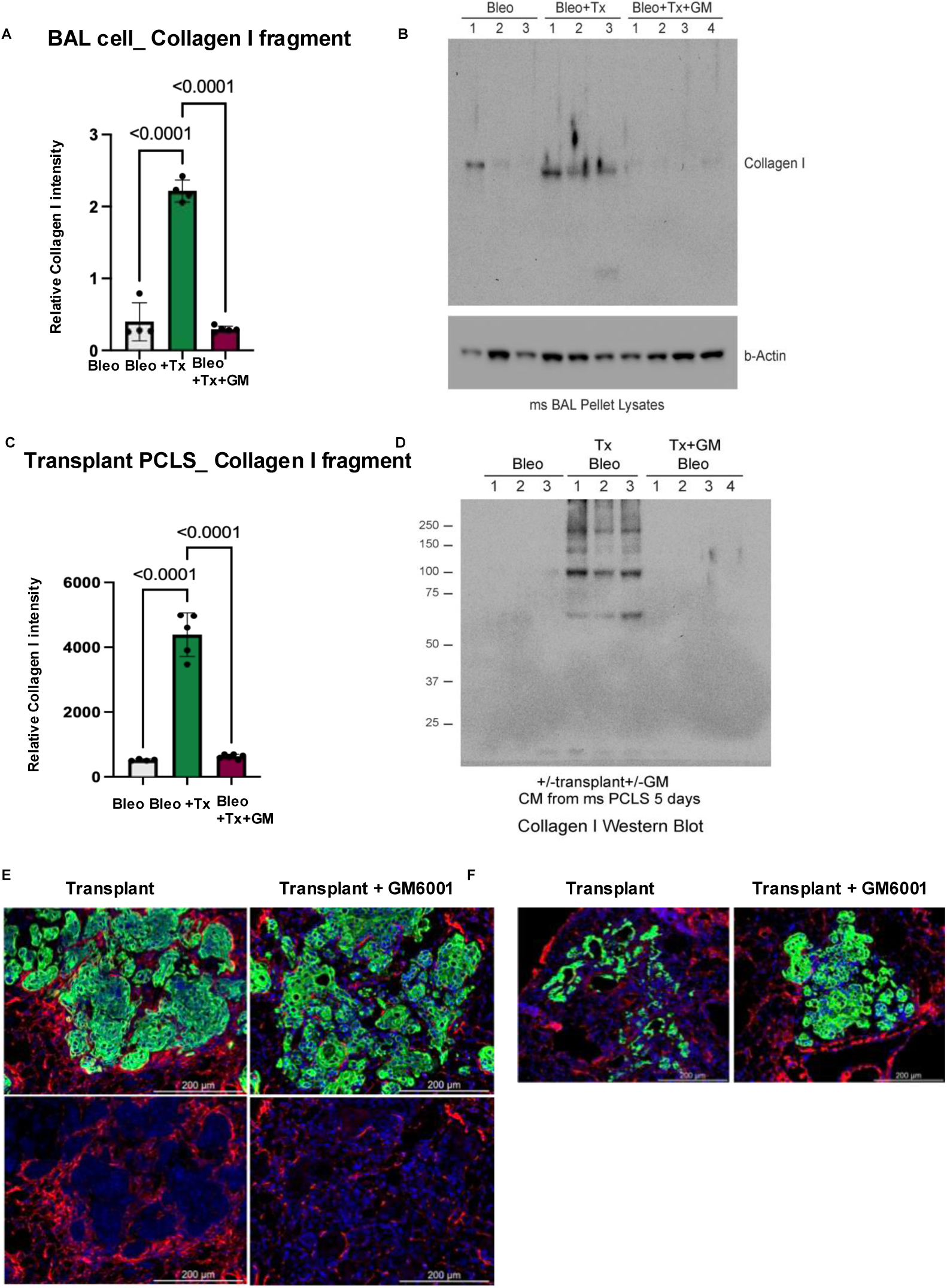
(**A**) Relative type I collagen intensity of (**B**) Western blot of degraded type I collagen in BAL cell lysates of bleomycin NSG mice transplanted with IPF BCs, treated with or without GM6001 (n = 3 – 4 biological replicates, 2 independent cohorts). (**C**) Relative type I collagen intensity of (**D**) Western blot of degraded type I collagen in concentrated conditioned media (CM) of PCLS from lungs of bleomycin NSG mice transplanted with IPF BCs, treated with or without GM6001 (n = 3 – 4 biological replicates, 2 independent cohorts). (**E**) Enlargement of images of region of engraftment showing collagen degradation area under IPF basal cells with or without GM6001 treatment. (**F**) Representative lung section images of microcyst-like formation of bleomycin NSG mice transplanted with IPF BCs that is abrogated by GM6001 treatment. Sections were stained by IF for human-specific KRT17 (green) and COL1A1 (red). Statistical significance was determined by Ordinary one-way ANOVA (A, C).

**Supplemental Figure 4:**
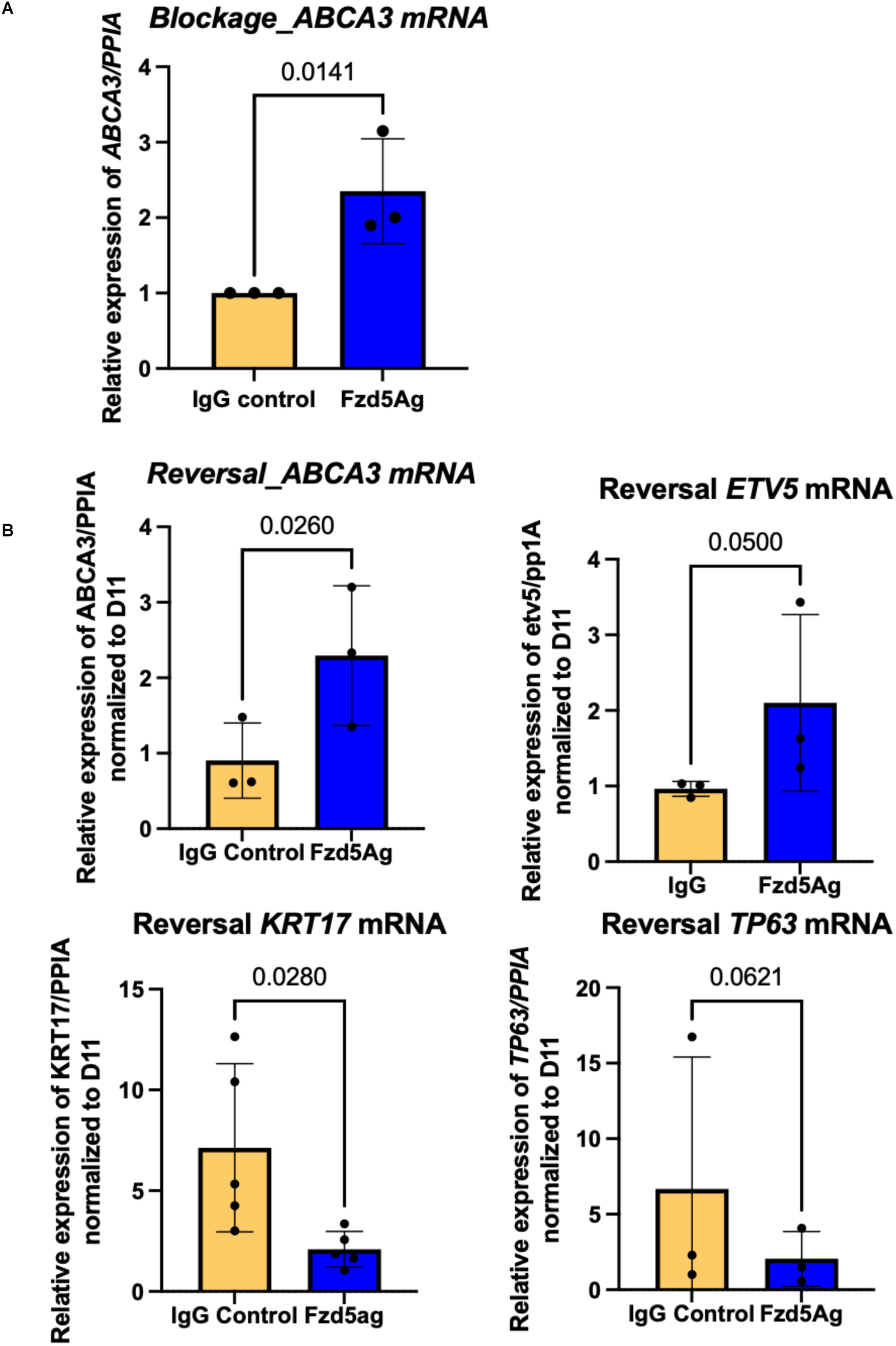
Additional cell fate markers expression after Fzd5Ag treatment of organoids. (A) *ABCA3* mRNA relative expression of epithelial cells (EPCAM+) sorted from organoids treated with IgG control or Frizzled-5 agonist (Fzd5Ag) from day 1 to day 14 of culture (n = 3). (B) *ABCA3*, *ETV5*, *KRT17*, and *TP63* mRNA relative expression of epithelial cells (EPCAM+) sorted from organoids treated with IgG control or Fzd5Ag from day 11 to day 19 of culture (n = 3). Statistical significance was determined by unpaired t-test (A) and paired t-test (B).

